# Continent-wide calibration of camera-trap metrics reveals low population densities in the European wildcat

**DOI:** 10.64898/2026.07.06.734798

**Authors:** Carolina Nogueira, Beatriz Alves, Stefano Anile, Javier Barona, Matteo Luca Bastianelli, Tamara Burgos, Marco Catello, Gonçalo Curveira-Santos, Francisco Díaz-Ruiz, Pau Federico, Christian Fiderer, Urša Fležar, Peter Gerngross, Jose María Gil-Sánchez, Maik Henrich, Javier Hernández-Hernández, Marco Heurich, Miha Krofel, Lea Maronde, Gonçalo Matias, Anna K. Moeller, Anja Molinari-Jobin, Anne Peters, Markus Port, Joe Premier, Filipe Rocha, Mariola Sánchez-Cerdá, Ferran Sayol, Marc Vilella, Emilio Virgós, Fridolin Zimmermann, José Jimenez, Pablo Ferreras, Pedro Monterroso

## Abstract

Effective conservation depends on demographic metrics that reliably reflect species status, particularly population abundance. For elusive species occurring at low densities, however, such metrics remain difficult to obtain. Spatial capture-recapture (SCR) models are the standardized approach for estimating density in marked populations, but their data requirements, especially the need for multiple spatial recaptures across individuals, often limit applicability in small or data-poor populations. This constraint has resulted in knowledge gaps for some of the most vulnerable species, undermining evidence-based conservation planning and management.

Using camera-trap data and SCR-derived density estimates from data-rich populations, we evaluated alternative, less data-demanding metrics and tested the hypothesis: Space to Event (STE), Mean Local Abundance (MLA), and Relative Abundance Index (RAI) exhibit predictable relationships with SCR-derived density; if supported, these metrics can reliably estimate density in populations where SCR models cannot be implemented. We applied this framework to the European wildcat (*Felis silvestris*), an elusive small felid with highly fragmented populations across Europe, for which density estimates are largely lacking despite growing conservation concern.

Across 21 study areas spanning most of the species’ range, our results indicate that European wildcats generally occur at lower densities than previously reported. SCR-derived estimates (n=10) averaged 10.32 ± 11.56 inds/100km^2^, while STE enabled density estimation in five additional data-poor areas (mean 5.52 ± 5.33 inds/100km^2^). STE showed a strong linear relationship with SCR-derived density (R^2^=0.98), supporting its use as a viable alternative when SCR is infeasible, although it tended to underestimate compared to SCR, especially at higher densities. In contrast, MLA and RAI showed weaker and non-linear relationships with SCR-derived density (R^2^=0.65), indicating substantially lower explanatory power and suggesting their estimates are more strongly influenced by confounding processes.

By explicitly calibrating alternative metrics across a wide density gradient throughout most of the species’ distribution, this study provides a transferable methodological framework for estimating density in low-density wildlife populations and the first continent-wide, standardized density assessment of a carnivore species. From a management perspective, our findings identify populations that may be most vulnerable, particularly those with the lowest densities, and highlight the need to prioritize absolute abundance monitoring.

## Introduction

Demographic parameters, particularly abundance and density, are cornerstones of population status assessments (Mills et al., 2025), as they are essential for quantifying population dynamics and extinction risk (Efford, 2004; Purvis et al., 2000; Stephens et al., 2015). Understanding the status and drivers of wildlife populations can help wildlife managers, policymakers, and the general public define and support conservation priorities, management decisions, and land use priorities (Heywood & Watson, 1995; Pellet & Schmidt, 2005; Stephens et al., 2015). Although closely related, abundance and density describe different aspects of population status. Abundance refers to the total number of individuals within a defined area, whereas density standardizes abundance by area, representing the number of individuals per unit area (Callaghan et al., 2024). Density is often considered more informative for ecological comparisons because it accounts for differences in spatial extent and allows comparisons among populations, habitats, or time periods (Royle et al., 2014). Although fundamental because they account for detection probability and therefore provide more accurate estimates, absolute abundance and density remain scarcer in comparison to estimates of relative abundance (i.e., a proxy of absolute abundance), as protocols for collecting, processing, and modeling the latter are typically simpler and less demanding (Callaghan et al., 2024). Therefore, robust monitoring programs must be defined to meet the analytical method’s assumptions and data requirements (Hayward et al., 2015; IUCN, 2012a; Popescu et al., 2016).

Spatial capture-recapture (SCR) is the modelling framework that currently provides the most robust estimates of population absolute abundance and density, as long as key assumptions are met, i.e., individuals can be uniquely identified, detected repeatedly, and sampled through a spatially explicit array of traps or detectors that captures the relationship between movement and detection (Crum et al., 2021; Morin et al., 2022). This framework extends classical capture-recapture models by considering the spatial heterogeneity in data and detection probability (Augustine et al., 2018; Royle et al., 2018; Sutherland et al., 2019). This approach has been suggested as particularly well-suited for detecting population density changes (Morin et al., 2022), and numerous studies have focused on optimizing sampling designs (e.g., designing new detector placement strategies) to meet the analytical requirements of SCR models (Dupont et al., 2021; Durbach et al., 2021; Efford & Fletcher, 2025; Jiménez et al., 2021; Palencia et al., 2024; Palmero et al., 2023). However, in practice, many camera-trap datasets are not originally designed for SCR analyses but for broader multi-species monitoring or general biodiversity surveys. As a result, obtaining sufficient individual identifications and spatial recaptures to fit SCR models can be challenging, particularly for elusive species or low-density populations (Brassine & Parker, 2015). To address such limitations, models integrating complementary data sources have been developed to refine parameter estimation in SCR models and obtain population estimates with higher precision (Jiménez et al., 2021; Ruprecht et al., 2021). Regardless of these advancements, their implementation still requires additional sources of data and advanced modelling and coding expertise, often unavailable to most ecologists, conservationists, or managers.

As a result, simple but reliable alternatives are required, such as relying on the more accessible unmarked animal data (O’Brien et al., 2003; O’Brien, 2011), which are often more compatible with multi-species inference goals in contrast to optimized species-specific sampling designs. For example, the Space to Event (STE) method (Moeller et al., 2018) relies on count data at individual camera stations and measures, through time-lapse sampling, the cumulative area sampled within the camera’s field of view before the first detection of the target species. The estimate obtained corresponds to the average density within the viewable area of the cameras, which can then be scaled to the area of interest (Moeller & Lukacs, 2021; Moeller et al., 2018). Even so, time-lapse sampling may prove challenging for rare or elusive species as it disregards detections occurring outside the defined intervals, significantly reducing the number of detections compared to passive motion sensor approaches.

Due to these potential limitations, abundance proxies may be the only viable option for some unmarked populations. For example, the mixture model with the Royle-Nichols parameterization (Royle & Nichols, 2003) is a hierarchical modelling approach that can be used to infer mean local abundance (MLA) from detection/non-detection data while modelling heterogeneity in detection probability. However, hierarchical models may still be challenging to fit in cases of small sample sizes, as is often the case with many elusive species. Given the limitations inherent to these more robust methods, the metric most widely used as a proxy of abundance derived from camera-trapping methods is the relative abundance index (RAI), which is calculated as the number of independent detections of the focal species per unit time, usually 100 trap days (O’Brien et al., 2003; O’Brien, 2011; Sollmann et al., 2013). Core assumptions of this analytical approach include constant detectability across space and time, and monotonic proportionality (i.e., a linear relationship) with true abundance (Sollmann et al., 2013). Although RAIs are simpler to estimate than true abundance (Link & Sauer, 1998; Pollock et al., 2002), these assumptions may not hold in most ecological contexts, leading to biases in the resulting estimates (Carbone et al., 2001; Carbone et al., 2002; Sollmann et al., 2013).

Although conservation priorities are guided by species-level assessments such as the International Union for the Conservation of Nature’s (IUCN) Red List of Threatened Species (Betts et al., 2020; de Lima et al., 2024), these global assessments often mask substantial knowledge gaps at local and regional scales (Palacio et al., 2023). Crucially, population abundance and its trends constitute core criteria underpinning IUCN Red List assessments, yet such information is frequently unavailable for many populations, thereby undermining robust evaluations of conservation status. As a result, data-deficient populations are likely to be disproportionately threatened, leading to distorted conservation priorities driven by uncertainty in extinction risk (Bland et al., 2015; Borgelt et al., 2022). This paradox is particularly pronounced in carnivores, whose biological traits – such as elusiveness and low population densities – hinder data collection and the estimation of reliable population metrics (Brassine & Parker, 2015; Johansson et al., 2020).

The European wildcat (*Felis silvestris*; hereafter wildcat) is an elusive, small felid typically occurring at low densities within fragmented populations (Gerngross et al., 2023; Gil-Sánchez et al., 2020; Lozano & Malo, 2012; Petisco et al., 2024). European wildcat individuals can be identified from camera-trap images based on their unique coat patterns (Anile et al., 2012). Although globally classified as Least Concern by the IUCN with an unknown population trend, population declines have been recorded in the Mediterranean region (Gerngross et al., 2023). The wildcat is considered Endangered in Portugal (Mathias et al., 2023), Vulnerable in Spain (IUCN, 2025), and Critically Endangered in Scotland (Breitenmoser et al., 2019). According to the IUCN Green Status assessment, the species is also recognized as Largely Depleted and not reaching the Functional state (Ambarli et al., 2025). In addition, knowledge gaps and the absence of reliable information about the current status, population sizes, and trends hinder the assessment of the wildcat conservation status over large parts of the species range (Gerngross et al., 2023). Given its ecological traits, phenotypic characteristics, and conservation needs, the wildcat emerges as an ideal model species to evaluate the reliability of alternative, less data-demanding metrics for estimating population density, while also benefiting from improved population size estimates in regions where sampling effort is suboptimal, thereby contributing directly to informed conservation decision-making. While genetic approaches remain indispensable for addressing hybridization (Kilshaw et al., 2015; Mattucci et al., 2019; Steyer et al., 2016), a critical concern for wildcat conservation, complementary camera trapping-based metrics offer a substantially cheaper means of obtaining population density estimates. The existence of camera trapping data already collected across multiple regions further provides a valuable foundation for testing such methods, even as species-optimized surveys remain a priority.

Using the wildcat as a model species, we tested the hypothesis: Space to Event (STE), Mean Local Abundance (MLA), and Relative Abundance Index (RAI) metrics exhibit proportional and monotonic relationships with population density estimated using benchmark SCR methods. Once calibrated, we expect that these metrics could reliably estimate or track density in data-poor populations where individual spatial recaptures prevent the implementation of SCR models. We tested this hypothesis by quantifying the relationship between SCR-derived density and alternative unmarked analytical approaches spanning a gradient of methodological complexity – STE, MLA, and RAI – obtained from camera trapping data across multiple wildcat populations spanning the species’ distribution. Subsequently, we applied the best-performing metrics to estimate population density of the wildcat across its distribution range, including populations where SCR-based estimation is not feasible or appropriate for sharing detectability and movement parameters under conditions of highly heterogeneous habitats. Through this hypothesis-driven framework, we aim to simultaneously contribute practical and scalable tools for assessing abundance from camera-trap data in low-density populations, and to provide the first continent-wide, standardized assessment of wildcat population status, thereby addressing a critical gap in the species’ conservation knowledge.

## Methods

### Study areas and data sources

The camera-trapping data were collected independently in 21 study areas from 8 European countries (Figure 1), either explicitly targeting the wildcat or where its detection was a bycatch of surveys focusing on other species (Mazzamuto et al., 2019). The 21 populations here span a variety of biomes, such as Mediterranean forests, woodlands, and scrublands, temperate broadleaf and mixed forests, and temperate conifer forests (Dinerstein et al., 2017; Figure 1). The Mediterranean biogeographical region is characterized by a warm climate with hot summers and mild winters (Condé & Richard, 2002; Rodriguez et al., 2005). In this biome, wildcat data were gathered from 13 populations across Portugal, Spain, Italy, and Albania. Temperate forests typically occur in the Atlantic and Continental biogeographical regions of Europe. The Atlantic region has a very long coastline influenced by the Atlantic Ocean and the North Sea. Hence, the climate is mild and humid. The Continental region is characterized by warm summers and cold winters (Condé & Richard, 2002). Wildcat populations were sampled in temperate forests across Portugal, Spain, Switzerland, Germany, Slovenia, and Austria. Finally, in the temperate conifer forests of the Eastern Italian Alps, located in the European Alpine biogeographical region, the annual and spatial distribution of rainfall is higher in the summer in the north, while the south is very dry in summer (Condé & Richard, 2002).

**Figure 1.**
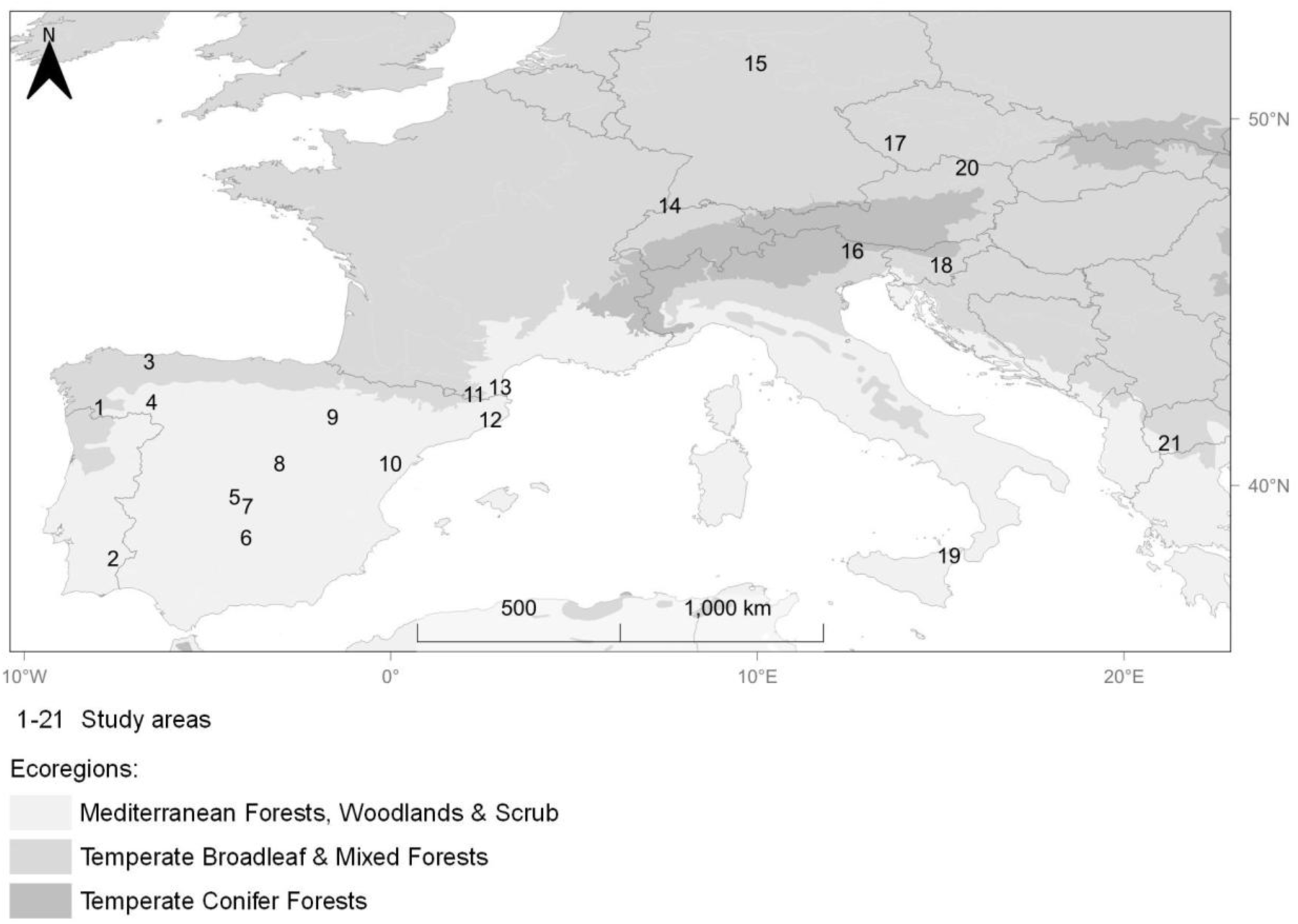
Geographic locations of the 21 study areas: 1 – Peneda-Gerês National Park, Portugal (PGNP); 2 – Guadiana Valley Natural Park, Portugal (GVNP); 3 – Muniellos Natural Reserve, Spain (MNR); 4 – Montesinho Natural Park, Portugal (MtNP); 5 – Cabañeros National Park, Spain (CNP); 6 – Sierra de Andújar Natural Park, Spain (SANP); 7 – Sierra de Picón, Spain (SP); 8 – Carabaña, Spain (Crb); 9 – Sierra de la Virgen, Spain (SV); 10 – Penyagolosa Natural Park, Spain (Pgl); 11 – Northern Catalunya, Spain (NC); 12 – Massís del Montseny Natural Park, Spain (MMNP); 13 – Northern Catalunya - Alta Garrotxa, Spain (NCAG); 14 – Northern Swiss Jura Mountains, Switzerland (NSJM); 15 – Melsunger Bergland, Germany (MB); 16 – Eastern Italian Alps, Italy (EIA); 17 – Bavarian Forest, Germany (BF); 18 – Northern Dinaric Mountains, Slovenia (NDM); 19 – Mount Etna, Italy (Etn); 20 – Wachau-Jauerling, Austria (WJ); 21 – Prespa National Park, Albania (PNP). Ecoregions according to Dinerstein et al. (2017).

Wildcat data were collected between 2009 and 2021, with the average study area inter-station distance varying between 371 and 2427 m (1118 ± 409, mean ± SD) (Appendix S1: Table S1). Sampling periods were standardized to ensure demographic closure (Karanth & Nichols, 2002; Royle et al., 2009), whereby maximum survey duration was set to 130-day primary sampling occasions as proposed for felid species occurring at low population densities (Brassine & Parker, 2015; Pollock, 1982). As a result, 2033 stations were deployed across 51 surveys. All seasons (i.e., summer, autumn, winter, and spring) were sampled in approximate frequency, ranging between 10-15 surveys/season (12.5 ± 1.8, mean ± SD). Survey effort varied between 5 and 210 cameras/survey (45.41 ± 49.31, mean ± SD), and mean survey length was 98.92 ± 38.23 (range: 30-130) (Appendix S1: Table S2).

Covariates were collected from survey methodological information to account for heterogeneity in detection probability within-studies due to their sampling designs, and included seasons, camera placement (on- or off-trail), camera model, flash type, number of cameras deployed per station, and presence of bait/lure (Anile & Devillard, 2016; Burton et al., 2015; O’Connell et al., 2011; O’Brien, 2011; Rovero & Zimmermann, 2016; Appendix S1: Tables S1 and S2).

### Estimation of density through SCR modelling

Individual identifications were performed by visual examination of coat patterns for the right, left, or both flanks, the latter in cases when individuals show both flanks in the same detection event (i.e., the set of photographs recorded during a single visit, captured by one or two cameras deployed per trapping station)(Johansson et al., 2020; Maronde et al., 2020)(Figure 2). Only records from putative wildcats were considered, based on coat characteristics as defined by Kitchener et al. (2005) and Ragni and Possenti (1996). Detection records with uncertain individual identity were discarded. Following Choo et al. (2020), additional procedures to prevent individual misidentification consisted of: (a) checking the photograph from each recapture against all previous photographs of the identified individual and other individuals; and (b) placing photographs side-by-side for comparison to ensure that no individual was assigned more than one identification code. All individual identifications were performed by the same observer, therefore reducing observer bias between study areas (Lioy et al., 2022). Dubious records were independently validated by a second observer. Individual tags were assigned using *Digikam* software (v.6.4.0).

**Figure 2.**
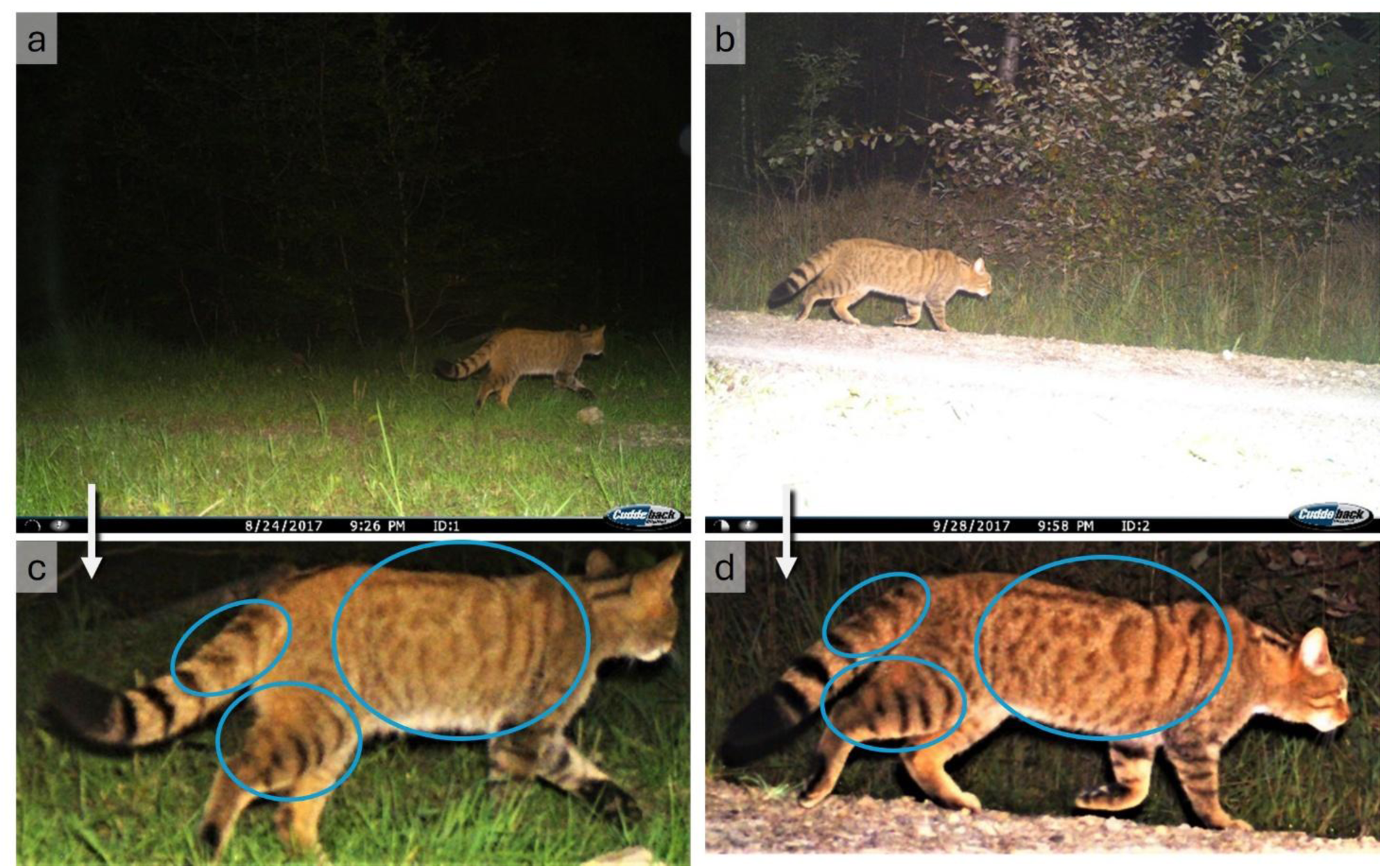
Individual MB_R2 captured during the night (white flash) on two different occasions and stations (a and b), and posterior image enhancement for comparison during the individual identification process (c and d).

All surveys with spatial recaptures were used to generate independent (single-season, closed-population) SCR density estimates in a Maximum Likelihood framework using the *oSCR* package v.0.42.0 (Sutherland et al., 2019) in *R* software v.4.0.3 (R Core Team, 2019). The discretized state space (SS) was defined using a buffer around the camera-trap locations, with a size informed by movement distances of detected individuals and set to 1.5 times the mean maximum distance moved (MMDM) in each study area (Matias et al., 2021). A resolution of 1000 m was used as an approximation of the minimum wildcat home ranges described in the literature (Oliveira et al., 2018; Ruiz-Villar et al., 2023). For surveys where the pooled MMDM was smaller than the defined grid resolution, a finer resolution (500 m) was used to maintain consistency between movement scale and state-space discretization. Sampling effort was accounted for by including a binary camera activity matrix, indicating whether at least one camera in each station was operational on each sampling occasion.

For each survey, we fitted a set of candidate models, including linear combinations of all detection covariates (i.e., temperate seasons, camera placement (on- or off-trail), camera model, flash type, number of cameras deployed per station, and presence of bait/lure) and a null model. Detection covariates were used to model the baseline detection rate (*p0*), i.e., detectability at the core of the theoretical home range. No covariates were used to parameterize the spatial scale parameter. We integrated the capture histories from each flank as different sessions in the same model, which was coerced to produce the same density estimate, thereby accounting for all individuals while avoiding double-counting identifications by different flanks (Chutipong et al., 2021; Matias et al., 2021; Meredith, 2018).

Models in each set were ranked and selected according to Akaike information criterion (AIC) (Akaike, 1973; Burnham & Anderson, 2002), where top-ranked models (i.e., the model with the lowest AIC) were considered most parsimonious in the set (Fuller et al., 2016). A full model including all parameters simultaneously was not feasible in most cases due to the small number of individuals detected per survey. Model selection was therefore necessary to identify the most supported parameter combination given the data available. Additionally, the coefficient of variation (CV) of each density estimate was generated to evaluate precision (Efford & Mowat, 2014; Jiménez et al., 2022; Palmero et al., 2023). In cases where the best model revealed a high CV (>0.6), an alternative model corresponding to a lower CV was chosen. Although CVs <0.30 have been suggested to indicate moderate precision (Dormann, 2013; Palmero et al., 2023), we took a more flexible approach with a <0.6 threshold, given the small number of individuals usually detected among surveys. When multiple surveys were carried out in the same study area, the survey for which density estimates revealed the lowest CV was selected. Although density can vary across surveys, the most accurate density estimate was considered the best representation of the area’s wildcat population. Unless otherwise stated, values are expressed as mean ± standard deviation (SD).

### Relationship between SCR and alternative metrics

We used all wildcat camera trapping records – marked and unmarked data – to estimate STE-derived density and alternative population size metrics (MLA and RAI) (Figure 3).

**Figure 3.**
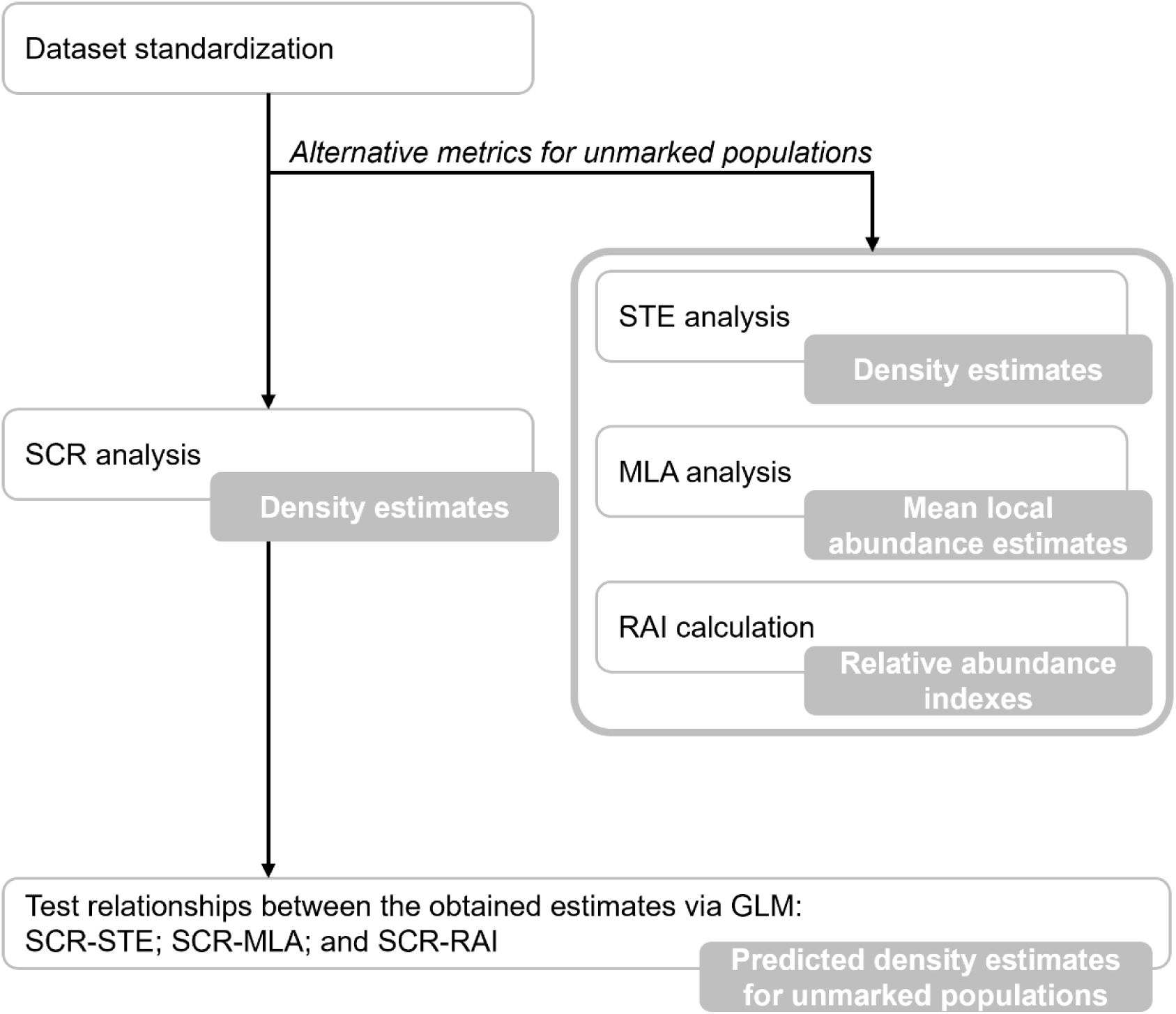
Data analysis workflow. SCR – Spatial Capture-Recapture; STE – Space to Event; MLA – Mean Local Abundance; RAI – Relative Abundance Index; GLM – Generalized Linear Model.

Space to Event estimates were generated with the *spaceNtime* R package (Moeller & Lukacs, 2021) using the camera model manufacturer specifications to calculate the viewshed area of each camera model (Appendix S1: Table S3). Near-instantaneous sampling occasions were defined with a frequency of 5 minutes and a sampling length of 10 seconds. Mean Local Abundance, defined as the number of animals associated with each camera station, was estimated with *occuRN* in unmarked v.1.0.1 (Fiske & Chandler, 2011). The same detection covariates and inclusion procedure as in the SCR approach were used to fit the MLA models. For each study area, the *ranef* function of the *unmarked* package was used to obtain the empirical Bayes estimates of abundance at each station, and the *bup* function to obtain the best unbiased predictor of abundance (Fiske & Chandler, 2011; MacKenzie et al., 2006; Royle, 2006). MLA estimates for each survey were calculated as the mean best unbiased predictor of abundance estimates across all camera stations. The RAI, defined as the number of independent wildcat detections per 100 trap days, was calculated for all study areas. The number of trap days of each survey was considered the cumulative number of days when at least one camera per station was active during the primary sampling period. Detection records were considered independent when the time between consecutive detections was >30 minutes (Rocha et al., 2021). Further details on the STE, MLA, and RAI analytical approaches can be found in Appendix S1: Section S2.

Weighted Generalized Linear Models (GLM) were fitted to assess the relationship between SCR-derived density and each corresponding STE, MLA, and RAI estimates, using the inverse of the uncertainty of the SCR-derived estimates as weight, in order to give greater influence to more precise estimates and reduce the contribution of less reliable ones, thereby improving the efficiency of the model fit. SCR-derived density estimates were set as the response variable, and the alternative metrics were set as independent covariates. The distribution family best suited to our data, among the gamma and the lognormal, was selected using graphical comparison of cumulative against fitted distributions, theoretical against empirical quantiles, and theoretical against empirical probabilities, and additionally by AIC comparison, using the *fitdistrplus* R package (Delignette-Muller & Dutang, 2015). Three link functions — identity, log, and inverse — were tested, and the model with the lowest AIC and best fit was selected. The models describing SCR-alternative metric relationships were evaluated using goodness-of-fit diagnostics, including R^2^, dispersion of residuals, and leave-one-out cross-validation. Values of R² < 0.5 were considered weak; 0.5 ≤ R² < 0.7 were considered moderate; and R² > 0.7 were considered strong in supporting the SCR-derived density estimates and alternative metric relationship (Moore, 1996). The *predict.glm* function of the *stats* R package (R Core Team, 2019) was then used to obtain predicted density estimates for study areas with poor data using SCR-alternative metric relationships with the most support.

## Results

We obtained 1,288 wildcat detections across all study areas and surveys, resulting in 143 individuals identified through the left flank and 151 through the right flank (Appendix S1: Table S4). Out of the total 51 surveys, only 11 (corresponding to 10 out of 21 study areas) yielded sufficient spatial recaptures to fit the SCR models and generate robust density estimates. Therefore, 40 surveys (78.43%) were not retained for the SCR analysis. Independent wildcat detections varied between 1 (PGNP in Portugal 2010, and NC in Spain 2016) and 141 (MB, Germany), averaging 25.25 ± 34.80 (mean ± SD) unmarked detections per survey (Appendix S1: Table S4).

### Estimation of density through SCR modelling

Among the 11 surveys retained for the SCR analysis, the average number of spatial recaptures was 1.58 ± 0.48, with the highest number achieved in WJ (Austria), with 2.5 spatial recaptures (i.e., mean number of unique trap detections per individual). The mean maximum distance moved averaged 2,354 ± 1,709 m and ranged from 681 m (NDM) to 6,640 m (WJ) (Appendix S1: Tables S5, S6, and S7).

SCR density estimates averaged 10.32 ± 11.56 individuals/100km^2^, ranging from 1.32 ± 0.47 individuals/100km^2^ in WJ to 34.18 ± 6.27 individuals/100km^2^ in MB (Table 1; Figure 5). Baseline detection probability (*p0*) was generally low, averaging 0.02 ± 0.02 and reaching a maximum of 0.09 ± 0.09 in WJ (2020, using Cuddeback Professional camera model deployed on trails; Appendix S1: Table S7). The spatial scale parameter averaged 1078 ± 679 m (range: 352-2631). (Appendix S1: Table S7).

**Table 1.**
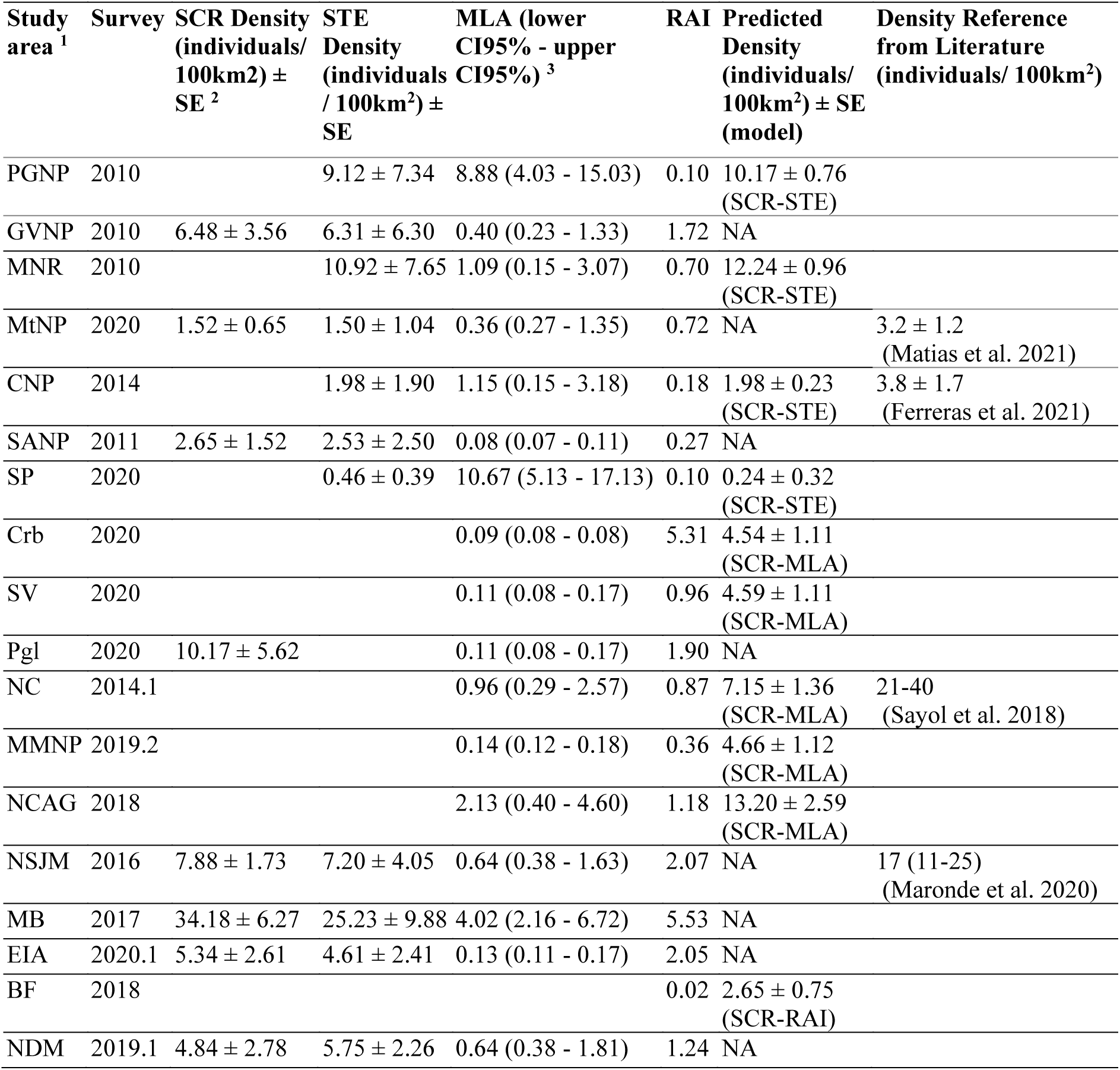
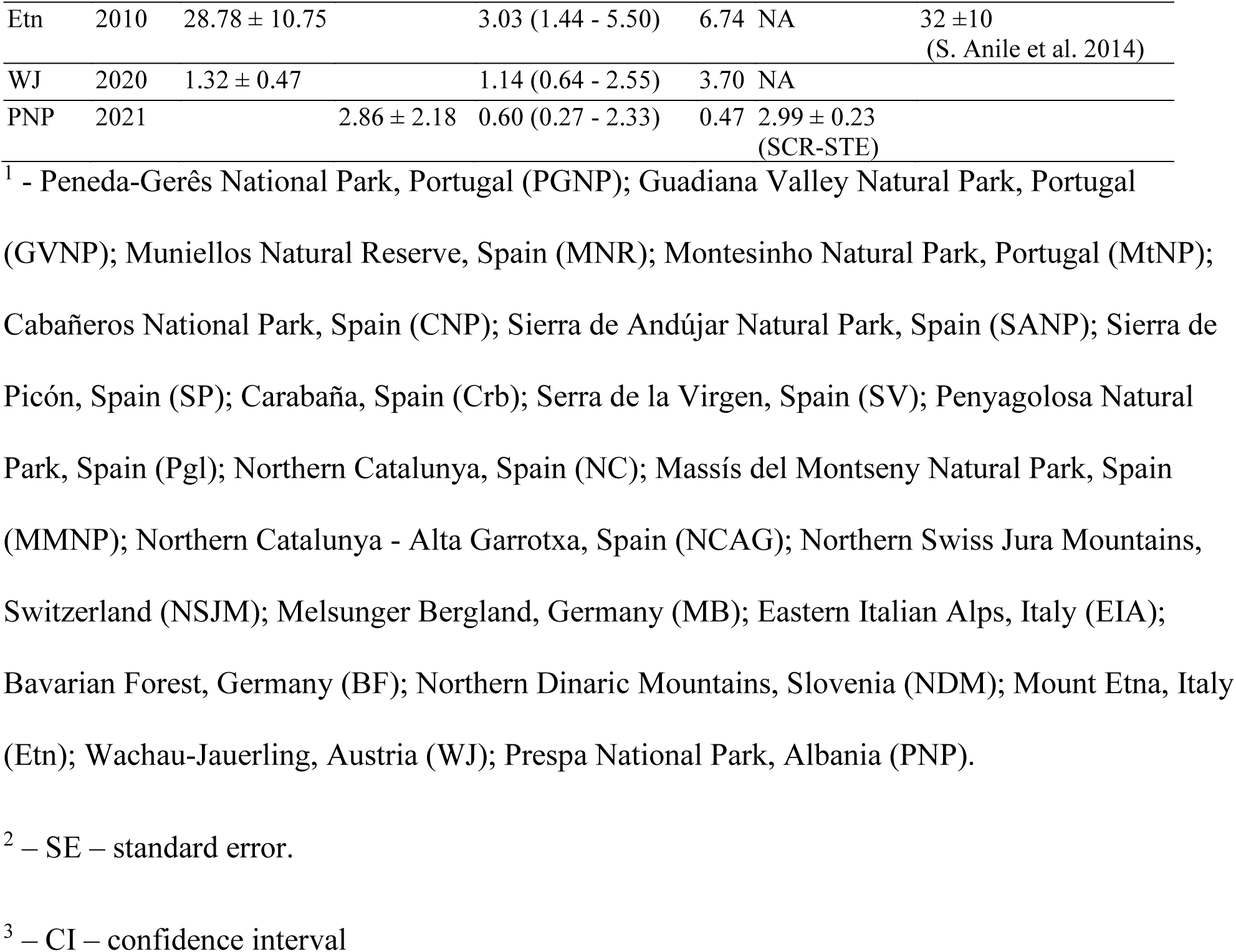
Estimates of spatial capture-recapture (SCR) derived density, Space to Event (STE) derived density, mean local abundance (MLA), relative abundance, and predicted density derived from the relationships of SCR to each alternative method - obtained for European wildcat populations. Previously published density estimates are provided as a reference. Study area, detections, and state space information can be found in Appendix S1: Table S1, Table S4, and Table S5, respectively.

### Relationship between SCR and alternative metrics

STE estimates were possible to be generated for 25 surveys (49%), corresponding to 12 out of the 21 study areas (57% study areas). MLA estimates were generated for 46 surveys (90%), corresponding to 20 (95%) study areas. RAI estimates were generated for all surveys (100%) (Table 1; Appendix S1: Tables S8, S9, and S10). GLM models between SCR and each alternative metric were generated for the 10 study areas with SCR-derived density estimates (Figure 4; Appendix S1: Table S11).

**Figure 4.**
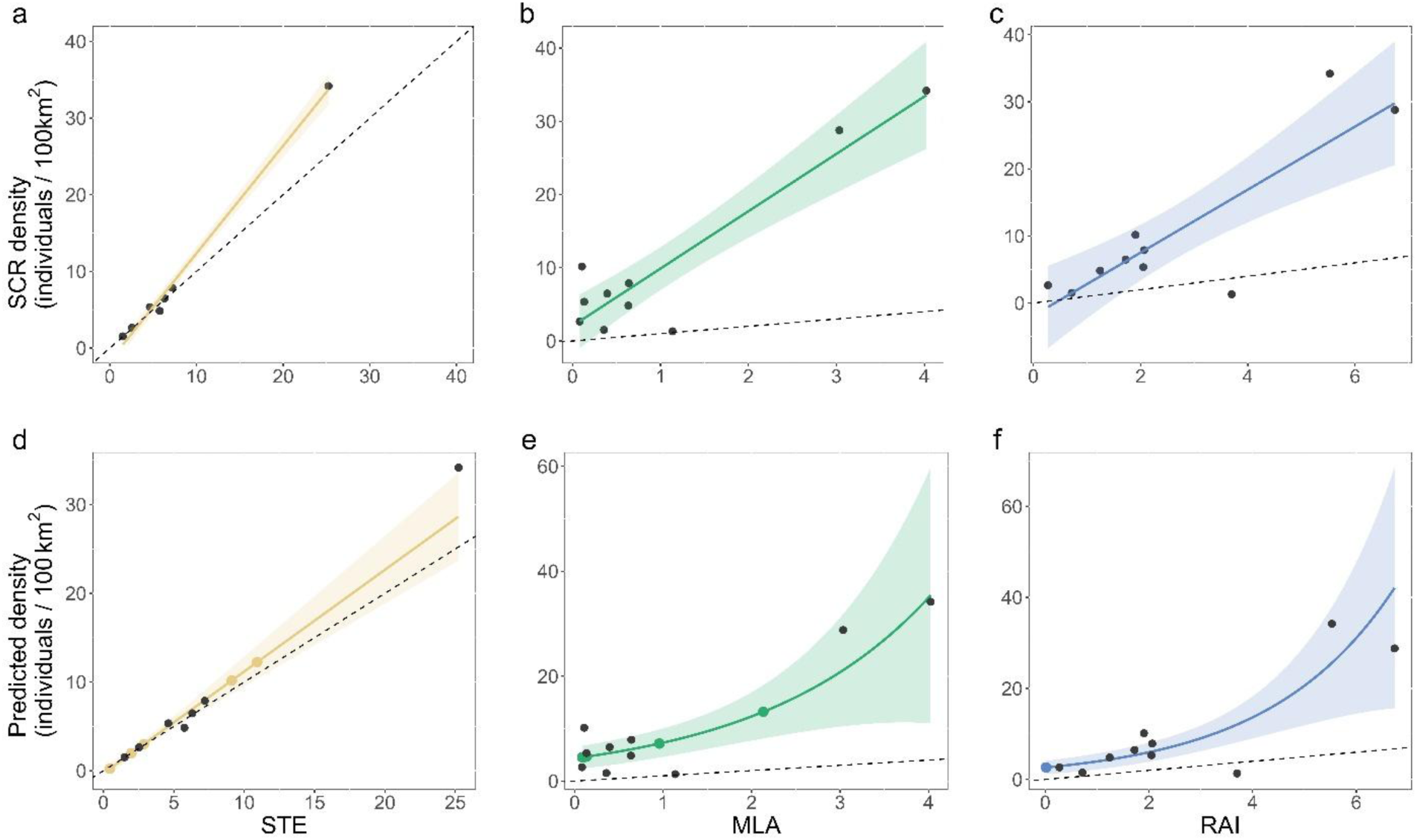
Relationships between each alternative metric and SCR (a, b, c) and resulting predicted density estimates (d - SCR-STE linear relation, e - SCR-MLA logarithmic relation, and f - SCR-RAI logarithmic relation). SCR estimates in grey; colored points for the study area estimates; colored lines for simulated values along the density range, and respective 95% confidence interval (ribbon); the dashed line is the identity line. Note: the scales differ between plots.

The Gamma error distribution provided the best fit for our data. The SCR–STE relationship was best fitted using an identity link, while the MLA–SCR and RAI–SCR relationships were better fitted using a log link (Appendix S1: Table S12). An R² = 0.98 was obtained for the SCR-STE model, indicating a strong relationship. Moderate support was found for SCR-MLA and SCR-RAI relationships, both with a R^2^ = 0.65. Goodness-of-fit diagnostics indicated the SCR-STE relationship had the highest support, followed by the SCR-MLA and SCR-RAI (Appendix S1: Table S13 and Figures S1 and S2). The resulting density prediction equation (Table 2 and Appendix S1: Table S14) allowed the generation of predicted density estimates where density was not obtained via SCR (Table 1, Appendix S1: Table S15, Figure 5).

**Table 2.** Density prediction and respective propagation of error equations to derive European wildcat population density estimates from Space to Event (STE), mean local abundance (MLA), and the relative abundance index (RAI).

| Model | Density prediction equation <sup>1</sup> | Propagation of error equation <sup>2, 3</sup> |
| --- | --- | --- |
| SCR-STE | **dpred** = 1.1472 * STE - 0.2912 | **SE** = $\sqrt{0.1259-STE \times 0.0610+STE^2 \times 0.0122}$<br>**Lower CI 95%** = dpred - 1.96 * SE<br>**Upper CI 95%** = dpred + 1.96 * SE |
| SCR-MLA | **dpred** = exp(0.5214 * MLA + 1.4687) | **SE** = **dpred** × $\sqrt{0.0629-MLA \times 0.0412+ MLA^2 \times 0.0140}$<br>**Lower CI 95%** = dpred - 1.96 * SE<br>**Upper CI 95%** = dpred + 1.96 * SE |
| SCR-RAI | **dpred** = exp(0.4114 * RAI + 0.9681) | **SE** = **dpred** × $\sqrt{0.0808-RAI \times 0.0362+ RAI^2 \times 0.0058}$<br>**Lower CI 95%** = dpred - 1.96 * SE<br>**Upper CI 95%** = dpred + 1.96 * SE |

<sup>1</sup> - dpred – predicted density derived from $\hat{\beta}_0 + \hat{\beta}_1 \times estimate$
<sup>2</sup> – SE – standard error derived from $\sqrt{Var(\beta_0)+2 \times dpred \times Cov(\beta_0, \beta_1)+ dpred^2 \times Var(\beta_1)}$
<sup>3</sup> – CI – confidence interval.

**Figure 5.**
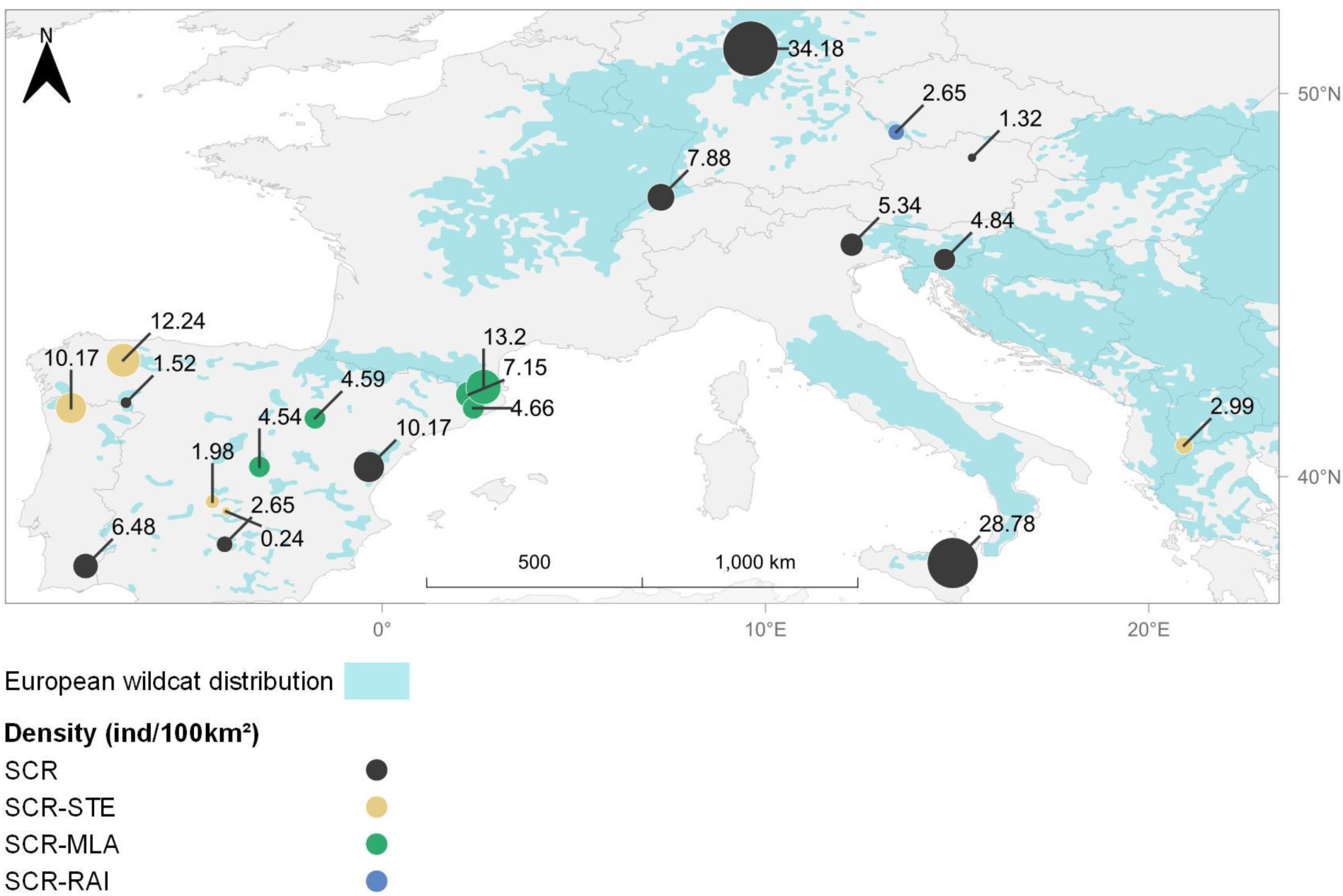
European wildcat densities across the species distribution range (adapted from the IUCN Red List of Threatened Species, version 2022-1, https://www.iucnredlist.org/). SCR - densities derived from spatial capture-recapture (grey); SCR-STE - densities predicted from Space to Event estimates (yellow); SCR-MLA - densities predicted from mean local abundance estimates (green); SCR-RAI - densities predicted from the relative abundance index (blue).

All our estimates were lower than previously estimated in the literature for the same populations (MtNP, CNP, NC, NSJM, and Etn; Table 1). Our study estimated an average wildcat density of 7.98 ± 8.64 individuals/100 km². This is lower than previously published estimates, which were on average 22.26 ± 12.39 individuals/100 km² (Anile et al., 2014; Dimitrijevic, 1980; Ferreras et al., 2021; Fonda et al., 2022; Gil-Sánchez et al., 2020; Heller, 1992; Kery et al., 2011; López-Martín et al., 2007; Maronde et al., 2020; Matias et al., 2021; Nussberger et al., 2023; Okarma & Olszańska, 2002; Ragni & Seminara, 2006; Sayol et al., 2018). Combining our results with published studies gives a range-wide average density of 14.76 ± 12.70 individuals/100 km² (range: 0.24–45.00; Figure 6).

**Figure 6.**
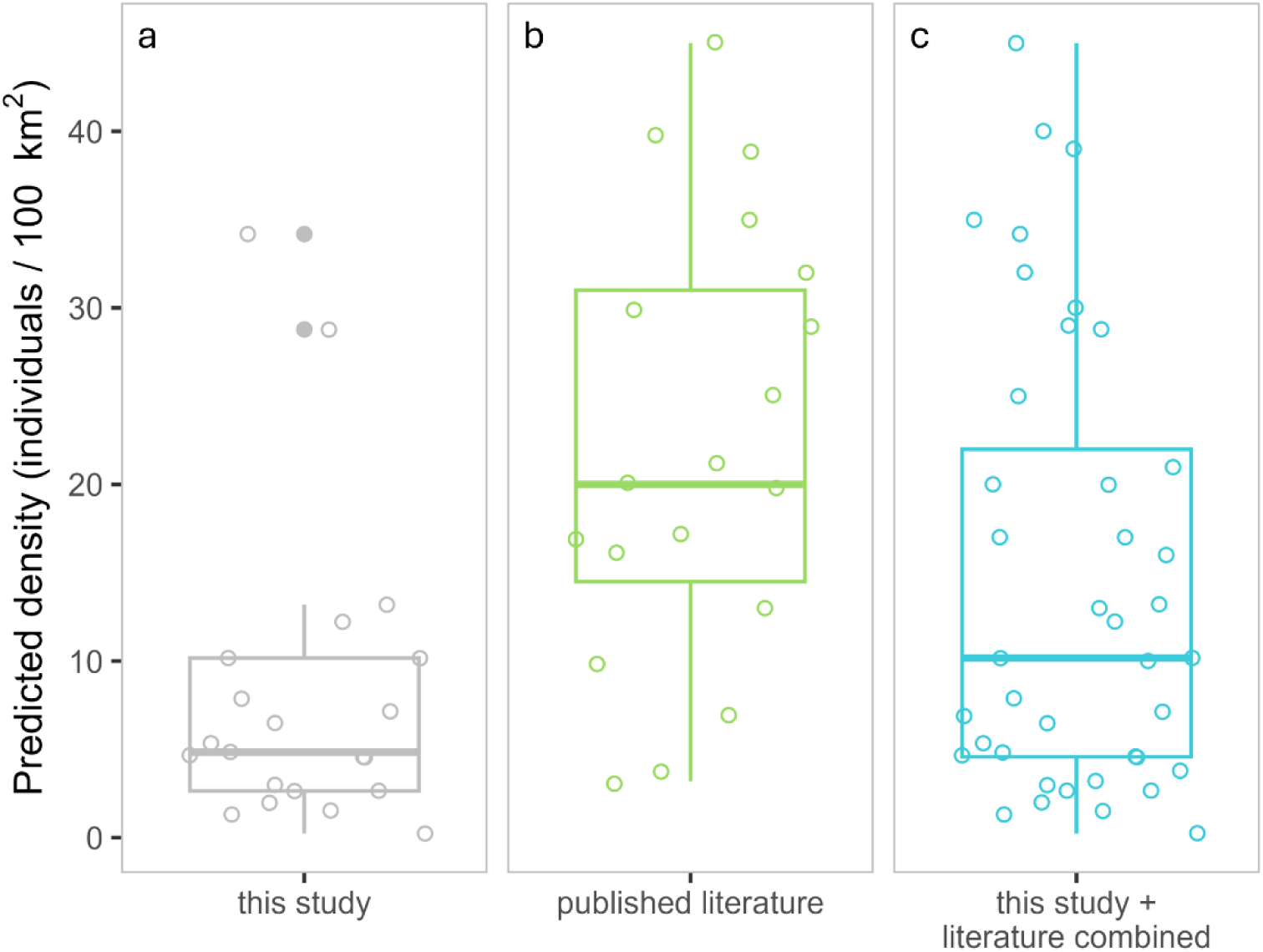
Distribution of European wildcat density estimates in: (a) this study, mean density of 7.98 ± 8.64 individuals/100km^2^; (b) published literature, mean density of 22.26 ± 12.39 individuals/100km^2^ (S. Anile et al. 2014; Dimitrijevic 1980; Ferreras et al. 2021; Fonda et al. 2022; Gil-Sánchez et al. 2020; Heller 1992; Kery et al. 2011; López-Martín et al. 2007; Maronde et al. 2020; Matias et al. 2021; Nussberger et al. 2023; Okarma and Olszańska 2002; Ragni and Seminara 2006; Sayol et al. 2018); (c) this study + literature combined, mean density of 14.76 ± 12.70 individuals/100km2 (range: 0.24 - 45.00).

## Discussion

Using the European wildcat as a model species, we described and quantified the relationship between benchmark Spatial Capture-Recapture estimates and three alternative, less data-demanding approaches – Space to Event, Mean Local Abundance, and Relative Abundance Index–spanning a gradient of methodological complexity. This allowed us to derive calibrated density functions that can be applied to additional wildcat populations. Importantly, albeit with varying precision levels, we produced density estimates for all 21 study areas, including populations lacking spatial recaptures, for which alternative metrics represented the only feasible option. To our knowledge, this study constitutes the first large-scale, continent-wide assessment of European wildcat density, delivering standardized estimates for 21 sites, exceeding the total number of studies previously available in the literature (n=19), and substantially improving the empirical basis for conservation status assessments across much of the species’ range, by providing the first estimates for several wildcat populations in Europe.

Among the methods considered, the relationship between SCR-derived density and all alternative metrics was positive and monotonic. However, only STE exhibited a strongly supported linear relationship with the benchmark estimates. The close agreement between SCR- and STE-derived density estimates supports our hypothesis and aligns with other works which suggest that STE is a reliable unmarked approach for estimating density in a solitary, low-density, and elusive species (Lyet et al., 2024; Punjabi et al., 2022). However, STE consistently slightly underestimated density compared to SCR (-1.39 ± 3.13 individuals/100 km², mean ± SD), especially at higher values. This pattern was previously attributed to the use of manufacturer-specified viewshed areas, i.e., a potential overestimation of the space in which an animal can be detected (Lyet et al., 2024). Future applications of STE could mitigate this bias by refining the effective camera viewshed, for instance, by estimating species-specific detection distances through field calibration experiments to truncate the effective detection area. Furthermore, although STE assumes random camera placement relative to animal movement (Moeller & Lukacs, 2021; Moeller et al., 2018), some camera locations in our study were deployed with the goal of increasing animal encounter probability, such as placement in trails, scent-marking sites or the use of attractants. Contrary to our results, these violations would be expected to cause an overestimation of density estimates, suggesting that this factor did not substantially influence our results.

In contrast, MLA and RAI only partly aligned with our hypothesis by showing log-linear relationships with SCR density. Our results indicate that only approximately 65% of the variation in these metrics reflects true differences in abundance, with the remaining variation likely driven by confounding factors such as heterogeneity in detectability across time, space and individuals (Kellner & Swihart, 2014; Sollmann et al., 2013), the absence of an explicit spatial component in the estimators (Amburgey et al., 2021), or unobserved ecological processes influencing movement and encounter rates (e.g., species-specific movement speed and grouping behaviors) (Gilbert et al., 2021). In addition, camera placement on trails, scent-marking sites or the use of attractants may introduce bias in unmarked approaches by potentially increasing encounter rates of the same individuals. Despite these limitations, both MLA and RAI proved useful proxies for abundance where more robust methods could not be applied, albeit with less reliability than STE.

The rapid global expansion of camera-trap monitoring for wildlife populations (Steenweg et al., 2017) has generated an increasing need for population metrics that can be derived from unmarked data, particularly for elusive and low-density species where capture–recapture approaches demand optimized and more complex sampling designs, as well as increased field and processing effort. While simple occupancy and count-based indices such as MLA and RAI have repeatedly been shown to suffer from high variance, low precision, and sensitivity to detectability and movement (Morin et al., 2022; Pettigrew et al., 2021; Sollmann et al., 2013; Vargas Soto et al., 2023; Wiegers et al., 2025), previous work has also suggested that they may retain value as calibrated proxies under restricted conditions (Carbone et al., 2001; Gil-Sánchez et al., 2020; Palencia et al., 2021; Parsons et al., 2017). Our study extends the literature by explicitly calibrating these metrics against SCR across a wide density gradient and across the species’ distribution, consistent with our hypothesis, but demonstrating that, in contrast to STE, their utility depends critically on species-specific and range-wide calibration rather than on point-estimate comparisons alone.

Consistent with previous assessments, we found that SCR- and STE-derived estimates could only be obtained for a subset of the study areas (47.62% and 57.14%, respectively), reflecting either suboptimal sampling designs or extremely low population densities that preclude reliable estimation with the most robust methods (Carbone et al., 2002). Conversely, MLA and RAI could be computed for the vast majority of the study areas, underscoring their relevance for data- deficient populations, provided uncertainty is properly propagated. These results highlight the need to optimize camera-trap survey design through adequate spatial coverage, sampling duration, and inter-camera spacing, to maximize the feasibility of SCR and STE approaches, particularly for rare species surveys (Dupont et al., 2021; Palmero, 2025; Palmero et al., 2023).

Our estimates revealed that most wildcat populations occur at low densities (<10 individuals/100km^2^), generally lower than those reported in earlier studies of the species (e.g., Dimitrijevic, 1980; Heller, 1992; Okarma & Olszańska, 2002; López-Martín et al., 2007; Ragni & Seminara, 2006). Many historical estimates were derived decades ago or relied on less robust methods, suggesting that the observed discrepancy may reflect a combination of population declines, overestimation by earlier approaches, or geographic bias toward higher-density populations, as is often the case with rare species (Jeliazkov et al., 2022). Despite its broad distribution, the wildcat fits the profile of a rare species characterized by consistently low densities, exemplifying the “rare species modelling paradox” whereby the species most in need of conservation attention are also the most difficult to monitor (Jeliazkov et al., 2022; Lomba et al., 2010; Rabinowitz, 1981). Indeed, the latest IUCN global assessment of the wildcat has recognized that knowledge gaps and the absence of robust abundance information have been a major limitation for assessing the species’ conservation status under the IUCN Red List and Green Status frameworks, particularly with respect to quantitative extinction-risk criteria (Collen et al., 2016; Gerngross et al., 2023).

This monitoring deficit is not unique to the wildcat but appears to characterize the genus *Felis* and other small felids more broadly. Our estimates are broadly consistent with the limited density data available for other small felids: they are in line with those reported for the black-footed cat (*Felis nigripes*; 3–8 ind/100km²; Sliwa et al., 2016), mainland leopard cat (*Prionailurus bengalensis*; 1.50-21.4 ind/100km^2^; Greenspan et al., 2025; Mohamed et al., 2013; Petersen et al., 2019; Rasphone et al., 2021), and marbled cat (*Pardofelis marmorata*; 3.8-19.6 ind/100km^2^; Hearn et al., 2016), lower than those of the jungle cat (*Felis chaus*; 20–150 ind/100km²; Gray et al., 2016) and the margay (*Leopardus wiedii*; 9.6-37.4 ind/100km^2^; Horn et al., 2020), and higher than the sand cat (*Felis margarita*; 1.12-2.9 ind/100km²; Abbadi, 1991; Gil-Sánchez et al., 2023). For the other small felids, including the Chinese mountain cat (*Felis bieti*) and the Afro-Asiatic wildcat (*Felis lybica*), no density estimates are available.

Our European wildcat density estimates were consistently lower than those reported in previous studies for the same populations (MtNP, CNP, NC, NSJM, and Etn; Table 1). Several factors may explain these discrepancies, including approaches to individual identifications (e.g., the number of observers and how conservative they are when assigning identifications) and differences in sampling duration. For instance, in NC, we restricted the sampling period to 130 days to comply with population closure assumptions (Brassine & Parker, 2015; Pollock, 1982), whereas Sayol et al. (2018) analyzed data collected over an entire year. Shorter sampling periods inevitably reduce the number of captures, which can result in lower abundance and density estimates. However, this does not necessarily reflect underestimation, as longer sampling periods risk violating the closure assumption (Dupont et al., 2019). Differences may also arise from the statistical modelling framework, such as the use of partial identity models in NSJM (Maronde et al., 2020) or the integration of occupancy data to refine SCR models (Ferreras et al., 2021). More generally, variation in SCR implementation, including monitoring duration (Gil-Sánchez et al., 2020; Kery et al., 2011), number of sampling occasions (Fonda et al., 2022), as well as population dynamics across studies conducted years apart, likely contributes to the observed differences among published density estimates (Fonda et al., 2022; Kery et al., 2011).

Marked spatial variation in density was also evident across regions. The WJ population in Austria exhibited the lowest density estimate, consistent with a slow recolonization following local extinction in the late 20^th^ century and limited population growth (Gerngross et al., 2021; Slotta-Bachmayr et al., 2016). At a broader scale, Mediterranean populations revealed slightly lower densities (average 6.84 ± 7.21 individuals/100km^2^), likely resulting from reduced prey availability in these ecosystems – the European rabbit (*Oryctolagus cuniculus*) –, declining habitat quality, and increased competition and intraguild predation in parts of the range where the Iberian lynx (*Lynx pardinus*) populations are being restored (Anile et al., 2019; Apostolico et al., 2016; Gil-Sánchez et al., 2020; López-Martín et al., 2007; Lozano et al., 2006; Monterroso et al., 2020; Moreno et al., 2008; Ruiz-Villar et al., 2023; Ruiz-Villar et al., 2026; Széles et al., 2018). These patterns are consistent with evidence of reduced genetic integrity of Mediterranean wildcat populations relative to other parts of the species’ continental range (Matias et al., 2022), whereby lower population densities, driven by prey scarcity, habitat degradation, and/or interspecific competition, likely reduce effective population sizes, limiting gene flow among fragmented subpopulations and increasing the risk of genetic drift and inbreeding (Frankham, 1995; Lande, 1988). Such demographic constraints may thus translate into the loss of genetic diversity, further aggravated by the increased hybridization rates with domestic cats, a well-documented threat to wildcat genetic integrity across Europe (Oliveira et al., 2008; Matias et al., 2022; Nussberger et al., 2014). This dynamic may mirror the relationship between population density and genetic diversity found in felids at the global scale (Azizan et al., 2023). Together, these findings highlight the importance of addressing the ecological drivers of reduced population density, particularly prey recovery and habitat restoration, not only for demographic reasons but also as a means of safeguarding the long-term genetic viability of wildcat populations across the Mediterranean range (Gil-Sánchez et al., 2020; Matias et al., 2022).

The main limitations of this study stem from: i) heterogeneity in sampling design among contributing projects; ii) from reliance on phenotypic identification of wildcats, without genetic confirmation; and iii) from non-independence across datasets, as the vast majority of the same data was used to estimate the alternative metrics. The reliance of phenotypic wildcats might have resulted in the inclusion of cryptic hybrids and exclusion of individuals that did not conform to typical phenotypic criteria (Devillard et al., 2014), although AI models appear to offer a robust solution to this task (Fargetta et al., 2025). We mitigated these limitations by standardizing sampling effort to the extent possible, explicitly modelling detection probability, and adopting conservative identification rules (i.e., only putative wildcats were considered, based on coat characteristics as defined by Kitchener et al. (2005) and Ragni and Possenti (1996); detection records with uncertain individual identity were discarded; and following Choo et al. (2020) guidelines to prevent individual misidentification). Nonetheless, poor image quality led to discrepancies between the number of detections and the number of individuals identified, which could have resulted in either under- or over-estimation of density, depending on whether exclusions were individual-specific or random (Creel et al., 2003; Johansson et al., 2020; Kodi et al., 2024; Sethi et al., 2016). However, despite yielding density estimates that are generally lower than those previously reported in the literature, including for some populations assessed here, the strong linear relationship between SCR- and STE-derived estimates, together with STE’s consistent tendency to slightly underestimate density relative to the benchmark, provides confidence that any potential bias in our SCR-derived density estimates would be systematic across all populations (i.e., affecting all estimates in a consistent, unidirectional manner), meaning that all estimates are subject to the same directional bias (either uniform underestimation or overestimation), such that the relative differences in density among populations remain valid and comparable. Finally, although SCR, MLA, RAI, and STE were derived from the same underlying sampling effort and are therefore not independent, by deriving the marked and unmarked datasets from the same field surveys, we minimized variation associated with sampling effort, habitat conditions, and animal behaviour, allowing differences among methods to be attributed primarily to their analytical frameworks and data requirements rather than to uncontrolled ecological or methodological variation.

Despite being the gold standard for density estimation of marked populations, SCR models remain challenging to fit to elusive species occurring at low population densities (Brassine & Parker, 2015). Reliable SCR estimation typically requires detecting more than 10 individuals per survey recaptured at least twice (Palmero et al., 2023; Schmidt et al., 2022), conditions rarely met for wildcats. In our study, the maximum number of individuals captured in a single SCR analysis was 17 (MB; Appendix S1: Table S5), with most surveys yielding fewer than five individuals and fewer than two spatial recaptures. Furthermore, some SCR estimates were obtained from surveys with relatively small camera arrays (i.e., <33 stations), which might further reduce precision (Dupont et al., 2021; Palmero, 2025). Under these conditions, alternative metrics to SCR become essential for informing the status of data-deficient and smaller populations, although we acknowledge their lower precision. We recommend that future monitoring efforts prioritize robust sampling designs that maximize detection of different individuals and spatial recaptures, thereby prioritizing SCR, when possible. Otherwise, STE stands as a more flexible and simple approach, but still a reliable abundance metric. While MLA and RAI can provide valuable information when more robust alternatives are infeasible, their use should be restricted to calibrated applications and considering uncertainty information (e.g., inverse-variance weighting). Given the tendency for discrepancy in density estimates observed in this study, we encourage future work to incorporate genetic confirmation, study designs optimized for detecting spatial recaptures, and statistical frameworks that explicitly model incomplete identity (e.g., SCR partial identity models). It should also be noted that our GLMs were built on only 10 data points, placing them at the lower threshold of what is typically considered sufficient for reliable regression analysis. We therefore encourage researchers to validate the calibrated density functions presented here against future SCR-based density estimates from additional wildcat populations as they become available.

The widespread occurrence of low-density populations contrasts sharply with the extensive geographic range of the wildcat across Europe (Gerngross et al., 2023). This disconnection underscores the need to move beyond distribution-based assessments alone towards multi-metric evaluations including viability and extinction risk at both population- and metapopulation-scales. Quantitative analysis of extinction risk forms the basis of Criterion E of the IUCN Red List, yet it remains rarely applied due to data limitations (Collen et al., 2016; IUCN, 2012b). Based on our findings, we suggest conservation efforts should prioritize monitoring populations at the lowest densities. By providing the first calibrated, continent-wide assessment of wildcat density based on benchmarked camera-trap methods, we offer both a transferable methodological framework for low-density species and a critical empirical foundation for reassessing conservation priorities across the species’ fragmented range.

## Acknowledgements

This paper was conceived and written within the collaborative EUROWILDCAT project (paper no. 007 of the EUROWILDCAT series; https://euromammals.org/eurowildcat/). The co-authors are grateful to all members for their support of the initiative. The EUROWILDCAT spatial database is hosted by Fondazione Edmund Mach. Fieldwork campaigns have been supported by LIFE Lynx LIFE16 NAT/SI/000634, Slovenian Research and Innovation Agency (grant J1-50013), Biodiversa+ (BIG_PICTURE project), Friends of the Earth Germany, HessenForst, and Project OAPN 352/2011. EUROwildcat was supported by the Regina-Bauer Stiftung, Prof Dr Riepe und Ellen Riepe-Brunnström Stiftung. BA’s work was supported by Fundação para a Ciência e a Tecnologia (FCT) through the PhD grant 2021.05061.BD, through the Portuguese State Budget and the European Social Fund (FSE/UE). TB was supported by a PhD fellowship (FPU17/04375) and a “Juan de la Cierva” postdoctoral contract (JDC2023-051114-I), funded by the Spanish Ministry of “Ciencia, Innovación y Universidades”. GC-S was funded by Fundacão para a Ciência e a Tecnologia (FCT) in the frame of Individual Call to Scientific Employment Stimulus Program (2023.08961.CEECIND/CP2845/CT0007). MH was supported by Ziel ETZ Free State of Bavaria – Czech Republic 2014–2020 (INTERREG V) – project number 184. FR is supported by National Funds through FCT-Fundação para a Ciência e a Tecnologia in the scope of the project UID/50027/2025-Research Network in Biodiversity and Evolutionary Biology. We thank all staff who supported the fieldwork campaigns. Particularly, Slovenia Forest Service personnel, Montseny Natural Park staff, and technicians from the Penyagolosa Natural Park. For the latter, work was carried out as part of the 1^st^ Spanish Wildcat Survey: special thanks to Juanjo Tur, Josep Puentes, and Miquel Ibañez. Finally, we are grateful to all volunteers, including local hunters, rangers, and students, who contributed to camera-trap revision and image annotation in NC, NCAG, EIA, and NDM.

## Author contributions

CN conceived and designed the study, compiled and standardized the data, performed the analyses, interpreted the results, wrote the first draft of the manuscript, and revised all versions of the manuscript. PM conceived and designed the study, provided data, contributed to the development of methods, and revised the manuscript. PF and JJ conceived and designed the study, provided data, and contributed to manuscript revision. SA, JB, MLB, TB, MC, FDR, PF, CF, UF, PG, JMG-S, MH, JH-H, MH, MK, LM, GM, AM-J, AP, MP, JP, MS-C, FS, MV, EV, and FZ provided data and contributed to manuscript revision. BA, GC-S, AM, and FR contributed to the development of methods and contributed to manuscript revision.

