## Appendix S1 for "Continent-wide calibration of camera-trap metrics reveals low population densities in the European wildcat"

**Preprint:** Submitted for publication

**Author names:** Carolina Nogueira, Beatriz Alves, Stefano Anile, Javier Barona, Matteo Bastianelli, Tamara Burgos, Marco Catello, Gonçalo Curveira-Santos, Francisco Díaz-Ruiz, Pau Federico, Christian Fiderer, Ursa Flezar, Peter Gerngross, Jose María Gil-Sánchez, Maik Henrich, Javier Hernández-Hernández, Marco Heurich, Miha Krofel, Lea Maronde, Gonçalo Matias, Anna Moeller, Anja Molinari-Jobin, Anne Peters, Markus Port, Joe Premier, Filipe Rocha, Mariola Sánchez-Cerdá, Ferran Sayol, Marc Vilella, Emilio Virgós, Fridolin Zimmermann, José Jimenez, Pablo Ferreras, Pedro Monterroso

### Section S1: Tables and Figures

Table S1 - Sampling design details for each of the 21 study areas. Surveys ranged between 2009 and 2021. Study areas: Peneda-Gerês National Park, Portugal (PGNP); Guadiana Valley Natural Park, Portugal (GVNP); Muniellos Natural Reserve, Spain (MNR); Montesinho Natural Park, Portugal (MtNP); Cabañeros National Park, Spain (CNP); Sierra de Andújar Natural Park, Spain (SANP); Sierra de Picón, Spain (SP); Carabaña, Spain (Crb); Serra de la Virgen, Spain (SV); Penyagolosa Natural Park, Spain (Pgl); Northern Catalunya, Spain (NC); Massis del Montseny Natural Park, Spain (MMNP); Northern Catalunya - Alta Garrotxa, Spain (NCAG); Northern Swiss Jura Mountains, Switzerland (NSJM); Melsunger Bergland, Germany (MB); Eastern Italian Alps, Italy (EIA); Bavarian Forest, Germany (BF); Northern Dinaric Mountains, Slovenia (NDM); Mount Etna, Italy (Etn); Wachau-Jauerling, Austria (WJ); Prespa National Park, Albania (PNP).

| Study area | Sampling period | Spatial design | Cameras per station | Inter-station distance (m) | Camera models | Flash type | Lures | Minimum convex polygon (km$^2$) |
|---|---|---|---|---|---|---|---|---|
| **MtNP** | 2020 | Grid | 1 | 1578 ± 636 | Cuddeback Model H-1453, Moultrie M-990i, Browning Strike Force HD Pro | Infrared | urine, valerian | 189 |
| **GVNP** | 2009-2010 | Grid | 1 | 1006 ± 327 | Leaf River IR5, Scout-Guard | Infrared | Urine, valerian | 145 |
| **PGNP** | 2010-2011 | Grid | 1 | 1028 ± 266 | Leaf River IR5, Scout-Guard | Infrared | Urine, valerian | 61 |
| **CNP** | 2009-2010 | Grid | 1 | 1100 ± 300 | Leaf River IR5, Scout-Guard | Infrared | Urine, valerian | 101 |
| **CNP** | 2012-2014 | Grid | 1 | 873 ± 355 | SG550V, SG570V HCO OutDoor Products, USA | Infrared | Urine, valerian | 103 |
| **MNR** | 2010-2011 | Grid | 1 | 1088 ± 293 | Leaf River IR5, Scout-Guard | Infrared | Urine, valerian | 105 |
| **SANP** | 2012 | Grid | 1 | 1092 ± 165 | Leaf River IR5, Scout-Guard | Infrared | Urine, valerian | 24 |
| **SANP** | 2011 | Opportunistic | 1 | 1233 ± 1531 | DLC Covert II | Infrared | Some stations (urine and pigeons) | 183 |
| **SP** | 2020 | Grid | 1 | 371 ± 283 | SR1-BK Spartan (HCO Outdoor Products, Georgia, USA | Infrared | Urine, valerian | 9 |

| | | | | | | | | |
|---|---|---|---|---|---|---|---|---|
| **Crb** | 2020 | Grid | 1 | 1461 ± 244 | BolyGuard | White | Urine | 19 |
| **Pgl** | 2020 | Grid | 1 | 1563 ± 207 | BolyGuard | White | Urine | 19 |
| **SV** | 2020 | Grid | 1 | 1471 ± 133 | BolyGuard | White | Urine | 16 |
| **MMNP** | 2018-2019 | Opportunistic | 1 | 1187 ± 326 | Browning Strike Force HD Pro, Cuddeback C | Both | None | 94 |
| **NC** | 2013-2016 | Grid | 1 | 833 ± 253 | Cuddeback C | Both | None | 14 |
| **NCAG** | 2017-2019 | Grid | 1 | 1516 ± 1037 | Cuddeback C | Both | None | 401 |
| **NSJM** | 2016 | Grid | 2 | 787 ± 215 | Cuddeback-Ambush, Cuddeback-Capture, Cuddeback-C1 | White | Valerian | 80 |
| **MB** | 2017 | Grid | 2 | 863 ± 207 | Cuddeback-Ambush, Cuddeback-C1 | White | None | 20 |
| **BF** | 2018-2019 | Grid | 1 | 1052 ± 163 | Cuddeback C2 | Infrared | None | 316 |
| **EIA** | 2017-2020 | Opportunistic | 1 | 2427 ± 2853 | UOVision, WildGame, Boskon, Ltl_Acorn | Infrared | None | 4197 |
| **EIA** | 2019-2020 | Grid | 1-2 | 751 ± 147 | Cuddeback, Secam, Browning, Moultrie | Both | None | 5 |
| **Etn** | 2010 | Line | 2 | 718 ± 272 | Sony-DSC-W55 | White | None | 11 |
| **NDM** | 2018-2019 | Opportunistic | 1-3 | 680 ± 1006 | Cuddeback X-Change Color Model 1279 | Both | Some stations (mix of catnip and beaver gland secretion in Vaseline) | 3221 |
| **WJ** | 2019-2020 | Opportunistic | 1 | 876 ± 809 | Cuddeback Professional, Cuddeback G, Cuddeback Ambush | White | Valerian | 19 |
| **PNP** | 2020-2021 | Grid | 1 | 1269 ± 302 | Cuddeback C | Infrared | None | 268 |

Table S2 - Effort summary: 2033 stations were deployed across 51 surveys. Survey effort varied between 5 and 210 cameras/survey (45.41 ± 49.31, mean ± SD), and mean survey length was 98.92 ± 38.23 (range: 30-130). (Survey – period of 130 days, at most, defined for the data analysis; Start/End – dates when sampling started/ended; #Stations – number of sampled stations; #Cameras – number of cameras deployed).

| Study area | Survey | Start | End | Sampling season | Sampling time (days) | #Stations | #Cameras |
|---|---|---|---|---|---|---|---|
| GVNP | 2009 | 30/07/2009 | 05/09/2009 | summer | 37 | 39 | 39 |
| | 2010 | 03/03/2010 | 05/04/2010 | spring | 33 | 39 | 39 |
| PGNP | 2010 | 05/10/2010 | 12/11/2010 | autumn | 38 | 35 | 35 |
| | 2011 | 05/04/2011 | 12/05/2011 | spring | 37 | 35 | 35 |
| MtNP | 2020 | 09/11/2019 | 18/03/2020 | autumn | 130 | 34 | 34 |
| CNP | 2009 | 24/09/2009 | 28/10/2009 | autumn | 34 | 44 | 44 |
| | 2010 | 26/01/2010 | 26/02/2010 | winter | 31 | 44 | 44 |
| | 2012 | 11/04/2012 | 19/06/2012 | spring | 69 | 60 | 60 |
| | 2013.1 | 20/02/2013 | 22/04/2013 | spring | 61 | 42 | 42 |
| | 2013.2 | 15/10/2013 | 19/11/2013 | autumn | 35 | 38 | 38 |
| | 2014 | 15/01/2014 | 23/04/2014 | winter | 98 | 40 | 40 |
| SP | 2020 | 01/02/2020 | 10/06/2020 | spring | 130 | 46 | 46 |
| MNR | 2010 | 24/08/2010 | 29/09/2010 | summer | 36 | 41 | 41 |
| | 2011 | 02/03/2011 | 04/04/2011 | spring | 33 | 42 | 42 |
| SANP | 2011 | 01/02/2011 | 11/06/2011 | spring | 130 | 28 | 28 |
| | 2012.1 | 29/02/2012 | 30/03/2012 | winter | 30 | 20 | 20 |
| | 2012.2 | 22/08/2012 | 16/10/2012 | autumn | 55 | 20 | 20 |
| NC | 2013 | 15/12/2013 | 24/04/2014 | winter | 130 | 7 | 7 |
| | 2014.1 | 25/04/2014 | 02/09/2014 | summer | 130 | 7 | 7 |
| | 2014.2 | 03/09/2014 | 31/12/2014 | autumn | 120 | 7 | 7 |
| | 2015 | 15/06/2015 | 23/10/2015 | summer | 130 | 6 | 6 |
| | 2016 | 24/10/2015 | 02/03/2016 | autumn | 130 | 6 | 6 |
| NCAG | 2017 | 12/08/2017 | 20/12/2017 | autumn | 130 | 5 | 5 |
| | 2018 | 21/12/2017 | 30/04/2018 | winter | 130 | 5 | 5 |
| | 2019.1 | 01/12/2018 | 01/04/2019 | winter | 121 | 12 | 12 |
| | 2019.2 | 02/04/2019 | 10/08/2019 | spring | 130 | 6 | 6 |
| MMNP | 2018 | 02/07/2018 | 09/11/2018 | summer | 130 | 18 | 18 |
| | 2019.1 | 10/11/2018 | 20/03/2019 | autumn | 130 | 18 | 18 |
| | 2019.2 | 22/05/2019 | 29/09/2019 | summer | 130 | 17 | 17 |
| Crb | 2020 | 19/11/2020 | 29/01/2021 | autumn | 71 | 12 | 12 |
| Pgl | 2020 | 15/06/2020 | 15/09/2020 | summer | 92 | 12 | 12 |
| SV | 2020 | 30/05/2020 | 30/08/2020 | summer | 92 | 12 | 12 |
| MB | 2017 | 26/06/2017 | 07/10/2017 | summer | 103 | 25 | 50 |
| BF | 2018 | 07/05/2018 | 14/09/2018 | summer | 130 | 108 | 108 |
| | 2019 | 11/04/2019 | 19/08/2019 | spring-summer | 130 | 106 | 106 |
| EIA | 2017 | 09/09/2017 | 17/01/2018 | autumn | 130 | 18 | 18 |
| | 2018 | 01/05/2018 | 08/09/2018 | summer | 130 | 20 | 20 |
| | 2019.1 | 19/07/2019 | 26/11/2019 | summer | 130 | 42 | 42 |
| | 2019.2 | 27/11/2019 | 05/04/2020 | winter | 130 | 44 | 44 |
| | 2020.1 | 06/04/2020 | 14/08/2020 | spring | 130 | 47 | 49 |
| | 2020.2 | 15/08/2020 | 23/12/2020 | autumn | 130 | 26 | 28 |

| | | | | | | | |
|---|---|---|---|---|---|---|---|
| Etn | 2010 | 14/05/2010 | 11/09/2010 | spring | 120 | 18 | 36 |
| WJ | 2019 | 13/11/2019 | 20/02/2020 | autumn | 99 | 8 | 8 |
| | 2020 | 21/02/2020 | 31/05/2020 | winter | 99 | 11 | 11 |
| NSJM | 2016 | 22/02/2016 | 30/04/2016 | winter | 68 | 64 | 128 |
| NDM | 2018 | 06/08/2018 | 14/12/2018 | autumn | 130 | 147 | 179 |
| | 2019.1 | 15/12/2018 | 24/04/2019 | winter | 130 | 122 | 151 |
| | 2019.2 | 20/08/2019 | 28/12/2019 | autumn | 130 | 155 | 210 |
| | 2020 | 29/12/2019 | 22/04/2020 | winter | 115 | 154 | 210 |
| PNP | 2020 | 27/08/2020 | 02/11/2020 | summer | 68 | 66 | 66 |
| | 2021 | 08/11/2020 | 18/03/2021 | winter | 130 | 55 | 55 |

Table S3 – Camera model manufacturer specifications and respective calculation of the viewshed area of each camera model used in the Space to Event analysis. The minimum area between the detection, viewshed and flash areas was used.

| **Camera model** | **Detection angle** | **Field of view** | **Detection distance (m)** | **Flash distance (m)** | **Detection area ($m^2$)** | **Viewshed area ($m^2$)** | **Flash area ($m^2$)** | **Minimum area ($m^2$)** |
|---|---|---|---|---|---|---|---|---|
| BolyMedia | 35 | 60 | 30 | 27 | 275 | 471 | 223 | 223 |
| Boskon Guard BG-526 | 67 | 56 | 25 | 13 | 365 | 305 | 83 | 83 |
| Browning/Prometheus Strike Force HD Pro/ BTC5HDP | 45 | 55 | 24 | 24 | 226 | 276 | 226 | 226 |
| Cuddeback | 28 | 40 | 12 | 30 | 35 | 50 | 220 | 35 |
| DLC Covert Scouting Cameras Covert II | NA | 40 | NA | 12 | NA | NA | 50 | 50 |
| Leaf River IR5 | 25 | NA | 16 | NA | 56 | NA | NA | 56 |
| Ltl Acorn Non panoramic model (Ltl-6511MG) | 55 | 52 | 15 | 25 | 108 | 102 | 283 | 102 |
| Moultrie M-40i | 40 | 40 | 24 | 24 | 201 | 201 | 201 | 201 |
| Moultrie M-990i | 40 | 40 | 18 | 18 | 113 | 113 | 113 | 113 |
| Moultrie S-50i | 40 | 40 | 30 | 30 | 314 | 314 | 314 | 314 |
| Scout-Guard SG550V | NA | 40 | 15 | 11 | NA | 79 | 42 | 42 |
| Scout-Guard SG570V | NA | 60 | 12 | 14 | NA | 75 | 103 | 75 |
| Secam | 52 | 52 | 20 | 20 | 181 | 181 | 181 | 181 |
| Spartan SR1-BK | 62 | 62 | 25 | 18 | 338 | 338 | 175 | 175 |
| Stealth Cam | 30 | NA | NA | 25 | NA | NA | 164 | 164 |
| UOVision | 52 | 48 | 15 | 15 | 102 | 94 | 94 | 94 |
| WildGame ST041N | 18 | 55 | 14 | 14 | 31 | 94 | 31 | 31 |

Table S4 – Number of independent detections and individual identifications per survey. Detections are considered independent when occurring at >30 minutes apart. Average spatial captures >1 are required to include the survey in SCR analysis. Non-identified detections are the number of encounters where the captured individual could not be identified due to poor quality of the photograph or uncertainty.

| Study area | Survey | Detections | Left-side identifications | Right-side identifications | Average left-side spatial captures | Average right-side spatial captures | Non-identified detections |
|---|---|---|---|---|---|---|---|
| GVNP | 2009 | 21 | 2 | 4 | 1 | 1 | 15 |
| | 2010 | 17 | 5 | 1 | 1.20 | 1 | 9 |
| PGNP | 2010 | 1 | 0 | 0 | 0 | 0 | 1 |
| | 2011 | 6 | 3 | 2 | 1 | 1 | 0 |
| MtNP | 2020 | 24 | 4 | 4 | 1.75 | 2.00 | 9 |
| CNP | 2009 | 3 | 0 | 1 | 0 | 1 | 2 |
| | 2010 | 7 | 0 | 0 | 0 | 0 | 7 |
| | 2012 | 10 | 2 | 2 | 1 | 1.50 | 5 |
| | 2013.1 | 3 | 0 | 1 | 0 | 1 | 2 |
| | 2013.2 | 2 | 1 | 1 | 1 | 1 | 1 |
| | 2014 | 7 | 2 | 2 | 1 | 1 | 6 |
| SP | 2020 | 6 | 1 | 1 | 2.00 | 1 | 3 |
| MNR | 2010 | 8 | 2 | 0 | 1 | 0 | 4 |
| | 2011 | 7 | 1 | 2 | 1 | 1 | 5 |
| SANP | 2012.1 | 0 | 0 | 0 | 0 | 0 | 0 |
| | 2012.2 | 2 | 0 | 1 | 0 | 1 | 1 |
| | 2011 | 8 | 2 | 2 | 1.50 | 1.50 | 3 |
| NC | 2013 | 7 | 0 | 2 | 0 | 1 | 5 |
| | 2014.1 | 8 | 2 | 0 | 1 | 0 | 4 |
| | 2014.2 | 7 | 1 | 0 | 3.00 | 0 | 1 |
| | 2015 | 4 | 0 | 2 | 0 | 1 | 2 |
| | 2016 | 1 | 0 | 0 | 0 | 0 | 1 |
| NCAG | 2017 | 7 | 1 | 1 | 1 | 1 | 5 |
| | 2018 | 7 | 0 | 0 | 0 | 0 | 7 |
| | 2019.1 | 10 | 5 | 2 | 1 | 1 | 2 |
| | 2019.2 | 13 | 4 | 4 | 1 | 1 | 0 |
| MMNP | 2018 | 10 | 3 | 1 | 1 | 1 | 5 |
| | 2019.1 | 12 | 2 | 1 | 1 | 1 | 7 |
| | 2019.2 | 6 | 1 | 2 | 1 | 1 | 4 |
| Crb | 2020 | 39 | 1 | 8 | 1 | 1 | 18 |
| Pgl | 2020 | 18 | 3 | 5 | 1 | 1.50 | 10 |
| SV | 2020 | 7 | 1 | 2 | 1 | 1 | 4 |
| MB | 2017 | 141 | 17 | 15 | 2.06 | 2.13 | 70 |
| BF | 2018 | 2 | 0 | 2 | 0 | 0 | 0 |
| | 2019 | 0 | 0 | 0 | 0 | 0 | 0 |
| EIA | 2017 | 2 | 0 | 0 | 0 | 0 | 2 |
| | 2018 | 2 | 0 | 0 | 0 | 0 | 2 |
| | 2019.1 | 47 | 1 | 1 | 1 | 1 | 45 |
| | 2019.2 | 82 | 8 | 7 | 1 | 1 | 58 |

| | | | | | | | |
|---|---|---|---|---|---|---|---|
| | 2020.1 | 79 | 7 | 2 | 1 | 1.50 | 67 |
| | 2020.2 | 27 | 2 | 2 | 1 | 1 | 23 |
| Etn | 2010 | 74 | 8 | 10 | 1.50 | 1.10 | 47 |
| WJ | 2019 | 30 | 3 | 3 | 1.67 | 1.33 | 9 |
| | 2020 | 36 | 2 | 3 | 2.50 | 1.67 | 16 |
| NSJM | 2016 | 86 | 10 | 12 | 2.40 | 2.08 | 23 |
| NDM | 2018 | 109 | 9 | 8 | 1 | 1 | 86 |
| | 2019.1 | 135 | 9 | 11 | 1.11 | 1.09 | 111 |
| | 2019.2 | 36 | 5 | 2 | 1 | 1 | 29 |
| | 2020 | 74 | 9 | 11 | 1.11 | 1 | 47 |
| PNP | 2020 | 11 | 1 | 4 | 1 | 1 | 4 |
| | 2021 | 27 | 3 | 4 | 1 | 1 | 20 |

Table S5 – Surveys included in the SCR approach due to the presence of spatial recaptures of the target species. (IDs - number of identifications; MMDM - mean maximum distance moved by the individuals identified by the left or right side; Buffer - distance defined according to equation 3; State-space - resulting area of inference for the density estimates; *NA* - Non-available MMDM value due to lack of spatial recaptures or identifications by one of the sides, therefore, data from the available side only was included.)

| **Study area** | **Survey** | **Left-side IDs** | **Right-side IDs** | **Average left-side spatial captures** | **Average right-side spatial captures** | **Left MMDM (m)** | **Right MMDM (m)** | **Pooled MMDM (m)** | **Buffer (m)** | **Resolution** | **State-space ($km^2$)** |
|---|---|---|---|---|---|---|---|---|---|---|---|
| GVNP | 2010 | 5 | 1 | 1.20 | 1 | 2062 | *NA* | 2062 | 3092 | 1000 | 158 |
| MtNP | 2020 | 4 | 4 | 1.75 | 2.00 | 2633 | 3720 | 3285 | 4928 | 1000 | 520 |
| CNP | 2012 | 2 | 2 | 1 | 1.50 | *NA* | 527 | 527 | 791 | 500 | 120 |
| SP | 2020 | 1 | 1 | 2.00 | 1 | 538 | *NA* | 538 | 806 | 500 | 17 |
| SANP | 2011 | 2 | 2 | 1.50 | 1.50 | 1115 | 1115 | 1115 | 1673 | 1000 | 250 |
| NC | 2014.2 | 1 | 0 | 3.00 | 0 | 1196 | *NA* | 1196 | 1795 | 1000 | 15 |
| Pgl | 2020 | 3 | 5 | 1 | 1.50 | *NA* | 1731 | 1731 | 2596 | 1000 | 69 |
| MB | 2017 | 17 | 15 | 2.06 | 2.13 | 1712 | 1730 | 1720 | 2580 | 1000 | 70 |
| EIA | 2020.1 | 7 | 2 | 1 | 1.50 | *NA* | 1298 | 1298 | 1947 | 1000 | 4595 |
| Etn | 2010 | 8 | 10 | 1.50 | 1.10 | 1495 | 3624 | 1921 | 2881 | 1000 | 62 |
| WJ | 2019 | 3 | 3 | 1.67 | 1.33 | 2831 | 5614 | 3758 | 5637 | 1000 | 241 |
| | 2020 | 2 | 3 | 2.50 | 1.67 | 6895 | 6386 | 6640 | 9961 | 1000 | 556 |
| NSJM | 2016 | 10 | 12 | 2.40 | 2.08 | 2654 | 3675 | 3091 | 4637 | 1000 | 270 |
| NDM | 2019.1 | 9 | 11 | 1.11 | 1.09 | 1289 | 74 | 681 | 1023 | 500 | 2578 |
| | 2020 | 9 | 11 | 1.11 | 1 | 176 | *NA* | 176 | 265 | 500 | 3175 |

Table S6 - SCR model AIC rank (1 model set for each survey). Model – parameterization of each model; AIC - Akaike Information Criterion; dAIC - difference between AIC score for the best model and the model being compared.

| Survey | Model | AIC | dAIC |
|---|---|---|---|
| GVNP, 2010 | ~1 | 125.34 | 0 |
| | p0~Trail | 127.26 | 1.92 |
| MtNP, 2020 | p0 ~ Model | 344.78 | 0 |
| | p0 ~ Trail + Model | 346.71 | 1.93 |
| | p0 ~ 1 | 348.57 | 3.79 |
| | p0 ~ Trail | 350.35 | 5.57 |
| SANP, 2011 | p0~Bait | 105.58 | 0 |
| | p0~Trail+Bait | 107.58 | 2 |
| | ~1 | 120.8 | 15.22 |
| | p0~Trail | 122.59 | 17.01 |
| NC, 2014.2 | ~1 | 65.22 | 0 |
| | p0~Model | 66.43 | 1.21 |
| | p0~White.flash | 66.43 | 1.21 |
| | p0~Trail | 66.84 | 1.62 |
| | p0~Trail+Model | 68.12 | 2.9 |
| | p0~Trail+White.flash | 68.12 | 2.9 |
| | p0~Trail+Model+ White.flash | 70.12 | 4.9 |
| MB, 2017 | p0 ~ Model | 1553.5 | 0 |
| | p0 ~ 1 | 1575.2 | 21.71 |
| EIA_2020.1 | p0~Trail | 183.84 | 0 |
| | p0~Trail+ Cams.per.Station | 185.31 | 1.47 |
| | p0~Trail+White.flash | 185.47 | 1.63 |
| | ~1 | 185.49 | 1.65 |
| | p0~Trail+White.flash+ Cams.per.Station | 186.93 | 3.09 |
| | p0~Cams.per.Station | 187.00 | 3.16 |
| | p0~White.flash | 187.21 | 3.37 |
| | p0~White.flash+ Cams.per.Station | 188.7 | 4.86 |
| | p0~Cams.per.Station+Model | 191.73 | 7.89 |
| | p0~Trail+ Cams.per.Station+Model | 193.43 | 9.59 |
| WJ, 2019 | p0~Model | 209.74 | 0 |
| | p0~Trail+Model | 211.65 | 1.91 |
| | ~1 | 215.68 | 5.94 |
| | p0~Trail | 217.12 | 7.38 |
| WJ, 2020 | p0~Trail+Model | 439.84 | 0 |
| | p0 ~ Model | 440.42 | 0.58 |
| | p0~Trail | 441.63 | 1.79 |
| | ~1 | 443.5 | 3.66 |
| NDM, 2019.1 | p0~White.flash | 410.83 | 0 |
| | p0~White.flash+ Cams.per.Station | 411.26 | 0.43 |
| | p0~Trail+White.flash | 412.83 | 2 |
| | p0~Trail+White.flash+ Cams.per.Station | 413.26 | 2.43 |
| | ~1 | 413.84 | 3.01 |
| | p0~ Cams.per.Station | 415.61 | 4.78 |
| | p0~Trail | 415.83 | 5 |
| | p0~Trail+ Cams.per.Station | 417.59 | 6.76 |

Table S7 - Parameters estimated for the best spatial capture–recapture (SCR) model selected for each study area. Covariate inclusion depended on the information available for each survey and its variability within the survey design. Best model – model selected according to lowest AIC and coefficient of variation <0.6; dAIC - Delta Akaike Information Criterion; SE – standard error; p0 - Baseline detection probability parameter estimate; (*) – removed from analysis due to coefficient of variation >0.6 and lack of alternative models; (**) - estimate for the study area selected according to the survey that generated lowest coefficient of variation; (***) - p0 varied depending on the detection variables, values presented below.

| **Study area** | **Survey** | **Model** | **dAIC** | **Density (individuals/ 100km$^2$)** | **Density SE** | **Density coefficient of variation** | **Sigma (m)** | **Sigma SE (m)** | **p0** | **p0 SE** |
|---|---|---|---|---|---|---|---|---|---|---|
| GVNP | 2010 | p0 ~ 1 | 0.00 | 6.48 | 3.56 | 0.55 | 728.06 | 204.42 | 0.021 | 0.016 |
| MtNP | 2020 | p0 ~ Model | 0.00 | 1.52 | 0.65 | 0.43 | 2630.77 | 458.37 | (***) | (***) |
| CNP | 2012 | p0 ~ 1 | 0.00 | 1.66 | 1.56 | 0.94 (*) | 421.76 | 274.44 | 0.017 | 0.021 |
| SP | 2020 | p0 ~ 1 | 0.00 | 4.26 | 4.47 | 1.05 (*) | 351.50 | 227.48 | 0.002 | 0.002 |
| SANP | 2011 | p0 ~ 1 | 15.22 | 2.65 | 1.52 | 0.57 (**) | 894.78 | 361.91 | 0.014 | 0.010 |
| NC | 2014.2 | p0 ~ 1 | 0.00 | 4.86 | 4.88 | 1.01 (*) | 854.49 | 285.94 | 0.042 | 0.035 |
| Pgl | 2020 | p0 ~ 1 | 0.00 | 10.17 | 5.62 | 0.55 | 538.54 | 164.61 | 0.024 | 0.021 |
| MB | 2017 | p0 ~ Model | 0.00 | 34.18 | 6.27 | 0.18 | 767.43 | 45.31 | (***) | (***) |
| EIA | 2020.1 | p0 ~ Trail | 0.00 | 5.34 | 2.61 | 0.49 | 659.22 | 160.77 | (***) | (***) |
| Etn | 2010 | p0 ~ 1 | 0.00 | 28.78 | 10.75 | 0.37 | 1340.34 | 351.04 | 0.004 | 0.002 |
| WJ | 2019 | p0 ~ Model | 0.00 | 1.82 | 0.85 | 0.47 | 1973.21 | 401.83 | (***) | (***) |
| | 2020 | p0 ~ Trail + Model | 0.00 | 1.32 | 0.47 | 0.36 (**) | 2197.29 | 366.75 | (***) | (***) |
| NSJM | 2016 | p0 ~ 1 | 0.00 | 7.88 | 1.73 | 0.22 | 1098.67 | 85.31 | 0.016 | 0.003 |
| NDM | 2019.1 | p0 ~ White flash | 0.00 | 4.84 | 2.78 | 0.57 | 642.13 | 349.48 | (***) | (***) |

*(***) Baseline detection probability parameters*

| **Study area** | **Survey** | **Best model** | **p0 parameters** | **p0 estimate** | **p0 SE** |
|---|---|---|---|---|---|
| MtNP | 2020 | p0 ~ Model | Prometheus BTC5HDP | 0.002 | 0.002 |
| | | | Cuddeback Ambush + Moultrie M 990i | 0.011 | 0.004 |
| | | | Moultrie M 990i | 0.000 | 0.000 |
| MB | 2017 | p0 ~ Model | Cuddeback C1 | 0.037 | 0.006 |
| | | | Cuddeback C1 + Cuddeback Ambush | 0.013 | 0.002 |
| | | | Cuddeback C1 + Cuddeback Attack | 0.012 | 0.006 |
| EIA | 2020.1 | p0 ~ Trail (*) | Off trail | 0.012 | 0.009 |
| | | | On trail | 0.003 | 0.002 |
| WJ | 2019 | p0 ~ Model | Cuddeback Ambush | 0.011 | 0.007 |
| | | | Cuddeback G | 0.014 | 0.012 |
| | | | Cuddeback Professional | 0.064 | 0.036 |

| | | | | | |
|---|---|---|---|---|---|
| | 2020 | p0 ~ Trail + Model | Cuddeback Professional + Off trail | 0.028 | 0.040 |
| | | | Cuddeback Professional + On trail | 0.088 | 0.087 |
| | | | Cuddeback G + Off trail | 0.004 | 0.004 |
| | | | Cuddeback G + On trail | 0.012 | 0.010 |
| | | | Cuddeback Ambush + Off trail | 0.009 | 0.008 |
| | | | Cuddeback Ambush + On trail | 0.031 | 0.012 |
| NDM | 2019.1 | p0 ~ White flash | Infrared flash | 0.002 | 0.001 |
| | | | White flash | 0.005 | 0.003 |

Table S8 - Space to event density estimates (number of individuals per 100km$^2$). CI 95% - confidence intervals with a level of confidence of 0.95; SE – Standard error.; Density varied between 0.46 ± 0.39 individuals/100km$^2$ in SP and 25.23 ± 9.88 in MB, averaging the 5.32 ± 4.88 individuals/100km$^2$.

| **Study area** | **Survey** | **Density (individuals/100km$^2$)** | **SE** | **CI 95%** | **Coefficient of Variation** |
|---|---|---|---|---|---|
| GVNP | 2009 | 8.37 | 6.75 | 2.14 - 33.79 | 0.86 |
| | 2010 | 6.31 | 6.30 | 1.24 - 32.2 | 1.00 |
| PGNP | 2010 | 9.12 | 7.34 | 2.34 - 36.72 | 0.85 |
| | 2011 | 9.21 | 7.52 | 2.32 - 37.63 | 0.86 |
| MtNP | 2020 | 1.50 | 1.04 | 0.45 - 5.2 | 0.78 |
| CNP | 2010 | 5.09 | 5.09 | 1.00 - 26.05 | 1.00 |
| | 2012 | 2.90 | 2.91 | 0.57 - 14.87 | 1.00 |
| | 2013.1 | 3.78 | 3.80 | 0.74 - 19.43 | 1.00 |
| | 2014 | 1.98 | 1.90 | 0.41 - 9.62 | 0.96 |
| SP | 2020 | 0.46 | 0.39 | 0.11 - 1.98 | 0.90 |
| MNR | 2010 | 10.92 | 7.65 | 3.16 - 37.7 | 0.70 |
| | 2011 | 5.41 | 5.42 | 1.06 - 27.71 | 1.00 |
| SANP | 2011 | 2.53 | 2.50 | 0.5 - 12.76 | 0.99 |
| MB | 2017 | 25.23 | 9.88 | 12.25 - 53.44 | 0.44 |
| EIA | 2019.1 | 2.78 | 1.92 | 0.84 - 9.58 | 0.76 |
| | 2019.2 | 3.97 | 2.08 | 1.60 - 10.73 | 0.69 |
| | 2020.1 | 4.61 | 2.41 | 1.77 - 12.13 | 0.52 |
| | 2020.2 | 3.10 | 2.70 | 0.73 - 13.64 | 0.91 |
| NSJM | 2016 | 7.20 | 4.05 | 2.63 - 20.32 | 0.63 |
| NDM | 2018 | 2.81 | 1.29 | 1.21 - 6.65 | 0.50 |
| | 2019.1 | 5.75 | 2.26 | 2.74 - 12.1 | 0.40 |
| | 2019.2 | 0.75 | 0.60 | 0.19 - 3.01 | 0.84 |
| | 2020 | 2.03 | 1.07 | 0.78 - 5.43 | 0.62 |
| PNP | 2020 | 4.42 | 3.77 | 1.07 - 18.98 | 0.90 |
| | 2021 | 2.86 | 2.18 | 0.77 - 10.84 | 0.80 |

Table S9 - Mean local abundance and respective confidence intervals for each survey. CI 95% - confidence intervals with a level of confidence of 0.95. Mean local abundance estimates (95% confidence interval) for the wildcat varied between 0.033 (0.026 - 0.026) in CNP and 14.250 (7.833 - 21.667) in NCAG, averaging 3.371 ± 4.604 individuals associated with each station.

| **Study area** | **Survey** | **Best model** | **Mean local abundance** | **CI 95%** |
|---|---|---|---|---|
| GVNP | 2009 | ~ Trail | 1.045 | 0.333 - 3.154 |
| | 2010 | ~ Trail | 0.395 | 0.231 - 1.333 |
| PGNP | 2010 | ~ 1 | 8.875 | 4.029 - 15.029 |
| | 2011 | ~ 1 | 0.206 | 0.083 - 1.083 |
| MtNP | 2020 | ~ 1 | 0.359 | 0.265 - 1.353 |
| CNP | 2009 | ~ 1 | 9.362 | 4.045 - 16.045 |
| | 2010 | ~ 1 | 0.896 | 0.136 - 3.136 |
| | 2012 | ~ 1 | 3.849 | 0.167 - 8.167 |
| | 2013.1 | ~ 1 | 9.918 | 4.071 - 16.071 |
| | 2013.2 | ~ 1 | 0.033 | 0.026 - 0.026 |
| | 2014 | ~ 1 | 1.145 | 0.150 - 3.175 |
| SP | 2020 | ~ 1 | 10.674 | 5.130 - 17.130 |
| MNR | 2010 | ~ Trail | 1.094 | 0.146 - 3.073 |
| | 2011 | ~ Model | 0.258 | 0.095 - 1.357 |
| SANP | 2012.2 | ~ 1 | 0.064 | 0.050 - 0.100 |
| | 2011 | ~ 1 | 0.083 | 0.071 - 0.107 |
| NC | 2013 | ~ Trail | 1.646 | 0.571 - 3.857 |
| | 2014.1 | ~ Trail | 0.957 | 0.286 - 2.571 |
| | 2014.2 | ~ 1 | 2.247 | 0.714 - 4.857 |
| | 2015 | ~ 1 | 12.093 | 5.667 – 19.000 |
| | 2016 | ~ 1 | 11.017 | 5.167 - 18.167 |
| NCAG | 2017 | ~ Model | 14.049 | 7.400 - 21.600 |
| | 2018 | ~ 1 | 2.132 | 0.400 - 4.600 |
| | 2019.1 | ~ Model | 13.107 | 6.750 - 20.500 |
| | 2019.2 | ~ Model | 14.250 | 7.833 - 21.667 |
| MMNP | 2018 | ~ 1 | 0.271 | 0.222 - 1.222 |
| | 2019.1 | ~ Model + Flash | 1.361 | 0.278 - 3.500 |
| | 2019.2 | ~ 1 | 0.135 | 0.118 - 0.176 |
| Crb | 2020 | ~ 1 | 0.086 | 0.083 - 0.083 |
| Pgl | 2020 | ~ 1 | 0.108 | 0.083 - 0.167 |
| SV | 2020 | ~ 1 | 0.108 | 0.083 - 0.167 |
| MB | 2017 | ~ 1 | 4.023 | 2.160 - 6.720 |
| EIA | 2017 | ~ 1 | 10.462 | 5.111 - 17.111 |
| | 2018 | ~ 1 | 10.294 | 4.100 - 17.100 |
| | 2019.1 | ~ Model | 0.138 | 0.095 - 0.190 |
| | 2020.1 | ~ Trail | 0.134 | 0.109 - 0.174 |
| | 2020.2 | ~ 1 | 0.039 | 0.038 - 0.038 |
| Etn | 2010 | ~ 1 | 3.034 | 1.444 - 5.500 |
| WJ | 2019 | ~ Model | 0.587 | 0.500 - 1.000 |
| | 2020 | ~ Trail + Model | 1.138 | 0.636 - 2.545 |
| NSJM | 2016 | ~ 1 | 0.640 | 0.375 - 1.625 |
| NDM | 2018 | ~ Cameras per station | 0.563 | 0.313 - 1.490 |
| | 2019.1 | ~ Trail + Cameras per station | 0.636 | 0.377 - 1.811 |

| | | | | |
|---|---|---|---|---|
| | 2019.2 | ~ 1 | 0.561 | 0.174 - 2.181 |
| | 2020 | ~ Trail + Model + Cameras per station | 0.394 | 0.214 - 1.234 |
| PNP | 2021 | ~ 1 | 0.601 | 0.273 - 2.327 |

Table S10 - Relative abundance indices. Wildcat relative abundance indexes varied between 0.016 in BF and 6.740 in Etn, averaging 1.223 ± 1.475.

| Study area | Survey | RAI |
|---|---|---|
| GVNP | 2009 | 1.786 |
| | 2010 | 1.721 |
| PGNP | 2010 | 0.099 |
| | 2011 | 0.572 |
| MtNP | 2020 | 0.715 |
| CNP | 2009 | 0.220 |
| | 2010 | 0.556 |
| | 2012 | 0.396 |
| | 2013.1 | 0.159 |
| | 2013.2 | 0.156 |
| | 2014 | 0.184 |
| SP | 2020 | 0.104 |
| MNR | 2010 | 0.703 |
| | 2011 | 0.580 |
| SANP | 2012.2 | 0.223 |
| | 2011 | 0.274 |
| NC | 2013 | 0.794 |
| | 2014.1 | 0.872 |
| | 2014.2 | 0.833 |
| | 2015 | 0.509 |
| | 2016 | 0.127 |
| NCAG | 2017 | 1.069 |
| | 2018 | 1.180 |
| | 2019.1 | 1.297 |
| | 2019.2 | 2.838 |
| MMNP | 2018 | 0.557 |
| | 2019.1 | 0.566 |
| | 2019.2 | 0.357 |
| Crb | 2020 | 5.306 |
| Pgl | 2020 | 1.903 |
| SV | 2020 | 0.964 |
| MB | 2017 | 5.529 |
| BF | 2018 | 0.016 |
| EIA | 2017 | 0.117 |
| | 2018 | 0.087 |
| | 2019.1 | 1.052 |
| | 2019.2 | 1.830 |
| | 2020.1 | 2.049 |
| | 2020.2 | 1.457 |
| Etn | 2010 | 6.740 |
| WJ | 2019 | 3.942 |
| | 2020 | 3.700 |
| NSJM | 2016 | 2.065 |
| NDM | 2018 | 0.689 |
| | 2019.1 | 1.242 |
| | 2019.2 | 0.228 |
| | 2020 | 0.713 |
| PNP | 2020 | 0.359 |

| | |
|---|---|
| 2021 | 0.468 |

Table S11 - Selected estimates for each study area, included in the glm models. Available SCR estimates were selected according to AIC and coefficient of variation. Only one survey per study area was selected to ensure independence of glm data points. Space to event (STE), mean local abundance (MLA), and relative abundance index (RAI) estimates were selected to match the same surveys as available for SCR.

| **Study area** | **Survey** | **oSCR Density (individuals/ 100km2)** | **oSCR SE** | **oSCR Coefficient of Variation** | **STE Density (individuals/ 100km2)** | **STE SE** | **STE Coefficient of Variation** | **MLA** | **MLA CI 95%** | **RAI** |
|---|---|---|---|---|---|---|---|---|---|---|
| GVNP | 2010 | 6.48 | 3.56 | 0.55 | 6.31 | 6.30 | 1.00 | 0.395 | 0.231 - 1.333 | 1.721 |
| MtNP | 2020 | 1.52 | 0.65 | 0.43 | 1.50 | 1.04 | 0.78 | 0.359 | 0.265 - 1.353 | 0.715 |
| SANP | 2011 | 2.65 | 1.52 | 0.57 | 2.53 | 2.50 | 0.99 | 0.083 | 0.071 - 0.107 | 0.274 |
| Pgl | 2020 | 10.17 | 5.62 | 0.55 | *NA* | *NA* | *NA* | 0.108 | 0.083 - 0.167 | 1.903 |
| MB | 2017 | 34.18 | 6.27 | 0.18 | 25.23 | 9.88 | 0.44 | 4.023 | 2.160 - 6.720 | 5.529 |
| EIA | 2020.1 | 5.34 | 2.61 | 0.49 | 4.61 | 2.41 | 0.52 | 0.134 | 0.109 - 0.174 | 2.049 |
| Etn | 2010 | 28.78 | 10.75 | 0.37 | *NA* | *NA* | *NA* | 3.034 | 1.444 - 5.500 | 6.74 |
| WJ | 2020 | 1.32 | 0.47 | 0.36 | *NA* | *NA* | *NA* | 1.138 | 0.636 - 2.545 | 3.7 |
| NSJM | 2016 | 7.88 | 1.73 | 0.22 | 7.20 | 4.05 | 0.63 | 0.64 | 0.375 - 1.625 | 2.065 |
| NDM | 2019.1 | 4.84 | 2.78 | 0.57 | 5.75 | 2.26 | 0.40 | 0.636 | 0.377 - 1.811 | 1.242 |

Table S12 – Link function model selection for each relationship ranked by AIC.

| Relationship | Model | Intercept | Estimate | Df | logLik | AICc | Delta |
|---|---|---|---|---|---|---|---|
| SCR-STE | Link = identity | -0.2912 | 1.147000 | 3 | -16.203 | 46.4 | 0.00 |
| | Link = log | 1.0050 | 0.104100 | 3 | -23.906 | 61.8 | 15.41 |
| | Link = inverse | 0.2529 | -0.008882 | 3 | -26.794 | 67.6 | 21.18 |
| SCR-MLA | Link = log | 1.4690 | 0.52140 | 3 | -81.842 | 173.7 | 0.00 |
| | Link = inverse | 0.1923 | -0.04135 | 3 | -82.506 | 175.0 | 1.33 |
| | Link = identity | 3.7700 | 5.83100 | 3 | -84.272 | 178.5 | 4.86 |
| SCR-RAI | Link = log | 0.9681 | 0.41140 | 3 | -82.002 | 174.0 | 0.00 |
| | Link = identity | 0.7670 | 3.69200 | 3 | -83.522 | 177.0 | 3.04 |
| | Link = inverse | 0.2099 | -0.02824 | 3 | -85.177 | 180.4 | 6.35 |

Table S13 – Leave-one-out cross-validation results.

| Model | Adjusted mean error (individuals/100km$^2$) |
|---|---|
| SCR-STE | 3.4 |
| SCR-MLA | 4.8 |
| SCR-RAI | 9.6 |

Table S14 - Coefficients, variance, and covariance of the best models used in the density prediction formulas. $\hat{\beta}_0$ – intercept coefficient; $\hat{\beta}_1$ – estimate coefficient; Var – Variance; Cov – covariance.

| **Model** | $\hat{\boldsymbol{\beta}}_0$ | $\hat{\boldsymbol{\beta}}_1$ | **Var($\boldsymbol{\beta}_0$)** | **Var($\boldsymbol{\beta}_1$)** | **Cov($\boldsymbol{\beta}_0, \boldsymbol{\beta}_1$)** |
|---|---|---|---|---|---|
| SCR-STE | -0.2912 | 1.1472 | 0.1259 | 0.0122 | -0.0305 |
| SCR-MLA | 1.4687 | 0.5214 | 0.0629 | 0.0140 | -0.0206 |
| SCR-RAI | 0.9681 | 0.4114 | 0.0808 | 0.0058 | -0.0181 |

Table S15 - Predicted density results (individuals/100km$^2$).

| Prediction Model | Study area | Year | Predicted density (individuals/100km2) | Standard error | Confidence interval (95%) |
|---|---|---|---|---|---|
| scr ~ ste | PGNP | 2010 | 10.17 | 0.76 | 8.68 – 11.67 |
| | SP | 2020 | 0.24 | 0.32 | 0.00[a] – 0.86 |
| | MNR | 2010 | 12.24 | 0.96 | 10.36 – 14.11 |
| | PNP | 2021 | 2.99 | 0.23 | 2.55 – 3.43 |
| | CNP | 2014 | 1.98 | 0.23 | 1.53 – 2.43 |
| scr ~ mla | NC | 2014.1 | 7.15 | 1.36 | 4.89 – 9.82 |
| | NCAG | 2018 | 13.20 | 2.59 | 8.12 – 18.28 |
| | MMNP | 2019.2 | 4.66 | 1.12 | 2.47 – 6.85 |
| | Crb | 2020 | 4.54 | 1.11 | 2.37 – 6.72 |
| | SV | 2020 | 4.59 | 1.11 | 2.46 – 6.78 |
| scr ~ rai | BF | 2018 | 2.65 | 0.75 | 1.18 – 4.12 |

[a] – Lower bound of the confidence interval was truncated to zero because negative values of density are biologically impossible.

Figure S1 – Plot of the fitted values (left) versus the residuals (right) of each relationship.

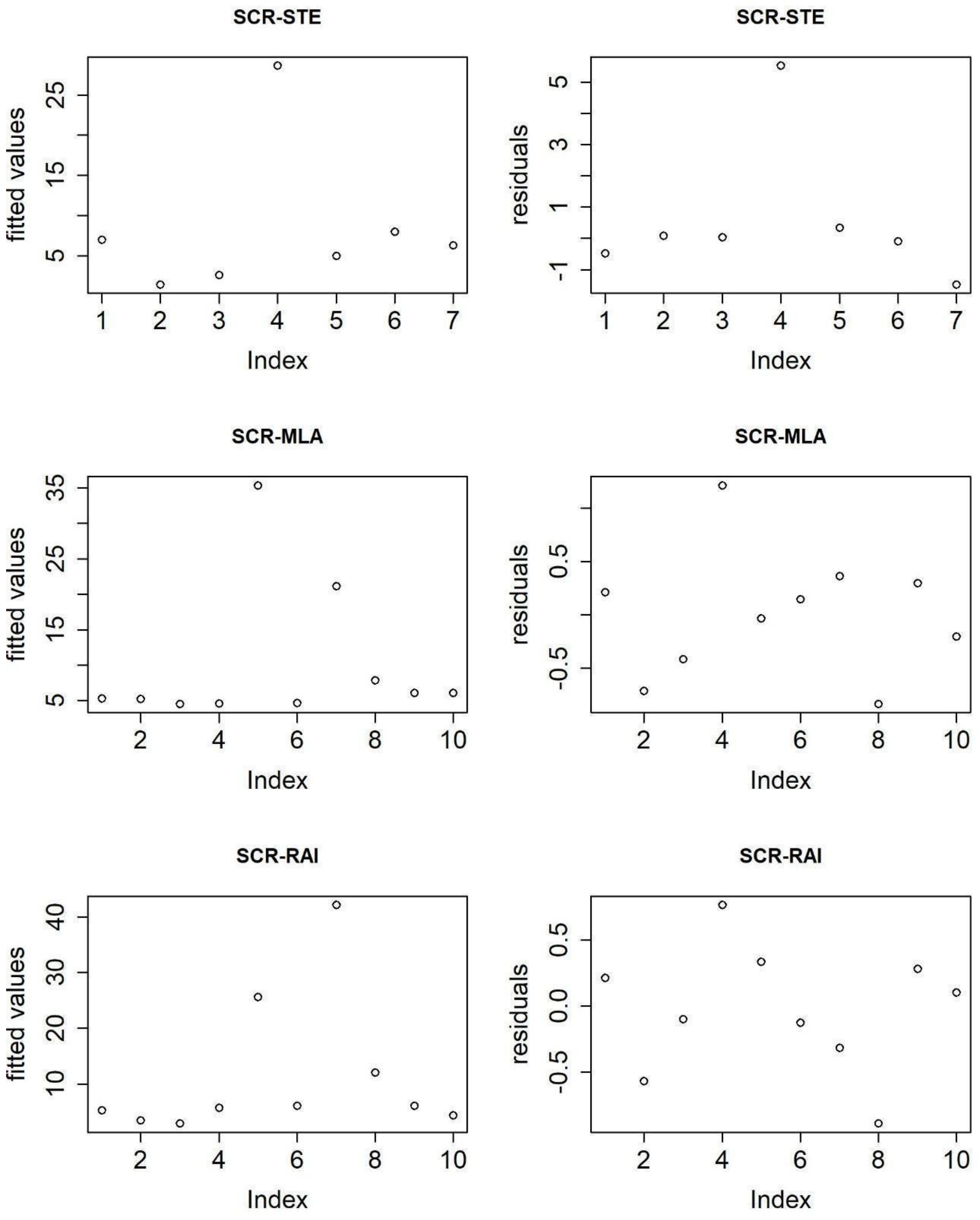

Figure S2 – Residuals QQ plot.

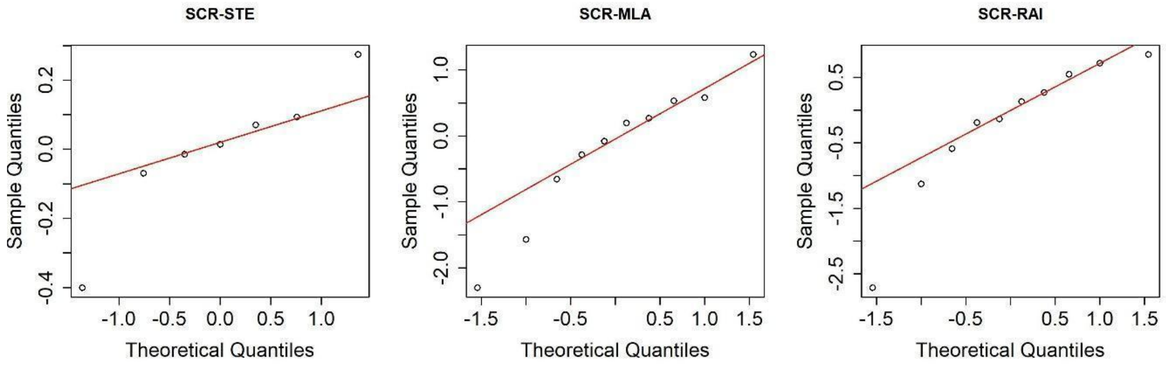

### Section S2: Alternative metrics applied in unmarked populations

#### Space to event (STE)

Space to event abundance estimates were generated with the *spaceNtime* R package (Moeller and Lukacs 2021) for the study areas where at least 25 camera stations were deployed. We used the camera model manufacturer's specifications to calculate the viewshed area of each camera model. Each station viewshed area was assumed to be constant across the whole survey length. For cases when more than one camera was deployed per station, station-level effort information was used, using functioning periods when at least one of the cameras was active, and assuming that both cameras were sampling overlapping areas.

Since we are working with species that are characterized by low detection rates, our goal was to achieve a trade-off between increasing sampling opportunity and not violating the instantaneous element of space to event models. As a result, instant sampling occasions were defined with a frequency of 5 minutes and a sampling length of 10 seconds. Additionally, to deal with the species' low detectability, the adaptation of motion-sensor data into instant sampling, and to increase the robustness and stability of results, for each survey, 5 versions of sampling occasions were created, each defined to start with a 0-, 1-, 2-, 3-, and 4-minute delay. As a result, most detections were able to be taken into account without an overestimation of abundance, by averaging the 5 estimates derived from each version.

A 100 $km^2$ study area was defined to facilitate comparison with the SCR estimates. For study areas where it was possible to estimate densities for more than one survey, the most accurate estimate was chosen based on the lowest coefficient of variation.

#### Mean local abundance (MLA)

Local abundances (i.e., the number of animals associated with each station of a certain study area) were estimated with *occuRN* in unmarked v.1.0.1 (Fiske & Chandler, 2011). Detection variables were used to fit the models in the same way as in the SCR approach. A MacKenzie and Bailey chi-square goodness of fit test was applied to the models, where p-values were obtained using a parametric bootstrap approach with 1000 samples (Mazerolle 2020). The p-value represents the proportion of the simulated test statistics greater than or equal to the observed test statistic (Mazerolle 2020). All p-values departing from either 0 or 1 were considered an acceptable fit (Kéry and Royle 2015). Overdispersion was tested by the c-hat value, which should be close to 1. Model sets validated by the goodness of fit were ranked and selected by AIC (Akaike 1973). For each survey, the model presenting the lowest AIC of the model set was selected, as considered to have the highest predictive performance in the set (Fuller et al. 2016). Whereas, for each study area, the model presenting the goodness of fit p-value closest to 0.5 was selected as the best fit and further used in the estimation of mean local abundances.

For each study area, the *ranef* function of the *unmarked* package was used to obtain the empirical Bayes estimates of abundance at each station. The *bup* function of the same package was applied to these estimates in order to obtain the best unbiased predictor of abundance for each station (Fiske and Chandler 2011; Royle 2006; MacKenzie et al. 2006). Mean local abundances for each study area were calculated as the mean best unbiased predictor of abundance estimates.

**Relative abundance index (RAI)**

The number of detections per 100 trap days, i.e., relative abundance index, was calculated for all study areas. The number of trap nights of each survey was considered the total number of days all cameras were active. Detection records were considered independent when the time between consecutive detections was >30 minutes (Rocha et al. 2021).

AICcmodavg: Model selection and multimodel inference based on (Q)AIC(c) R package version 2.3-1.

Moeller, Anna K., and Paul M. Lukacs. 2021. "spaceNtime: an R package for estimating abundance of unmarked animals using camera-trap photographs." *Mammalian Biology*. https://doi.org/10.1007/s42991-021-00181-8.

Rocha, M., A. Serronha, M. Rodrigues, P. C. Alves, and P. Monterroso. 2021. "Comfort over safety: thermoregulation overshadows predation risk effects in the activity of a keystone prey." *Journal of Zoology* n/a (n/a). https://doi.org/https://doi.org/10.1111/jzo.12947.

Royle, J. Andrew. 2006. "Site Occupancy Models with Heterogeneous Detection Probabilities." *Biometrics* 62 (1): 97-102.
